# Scaffold-mediated miRNA-155 inhibition promotes regenerative macrophage polarisation leading to anti-inflammatory, angiogenic and neurogenic responses for wound healing

**DOI:** 10.1101/2025.10.16.682055

**Authors:** Juan Carlos Palomeque Chávez, Marko Dobricic, Matthew McGrath, Cian O’Connor, Tara K. McGuire, Jack Maughan, Adrian Dervan, James E. Dixon, Cathal J. Kearney, Shane Browne, Fergal J. O’Brien

**Author notes:** **Corresponding Author:**, Address: 123 St. Stephen’s Green, Dublin 2, D02YN77, Ireland.

## Abstract

Chronic wounds represent a significant clinical challenge due to persistent inflammation and impaired nerve regeneration, both of which delay healing. Conventional treatments often yield limited success and inconsistent outcomes, especially in complex and recurring wounds. Combinatorial strategies that integrate biomaterial scaffolds with gene delivery offer a promising approach to promote tissue repair. MicroRNAs (miRNAs), particularly miRNA-155, have emerged as crucial regulators of wound healing. MiRNA-155 is highly expressed in inflammatory conditions and plays a key role in macrophage activation, polarisation, and nerve regeneration. In this context, this study introduces a miRNA-155 inhibitor-activated scaffold designed to modulate the chronic wound environment by inhibiting miRNA-155. MiRNA-155 inhibitor complexed GET nanoparticles were characterised and incorporated into collagen-glycosaminoglycan (CG) scaffolds. These supported dermal fibroblast and endothelial cell growth while enabling controlled inhibitor release. Scaffold-mediated miRNA-155 inhibition in both non-polarised (M0) and pro-inflammatory (M1) macrophages promoted anti-inflammatory (M2) polarisation. Macrophage secretome analysis showed reduced inflammatory cytokines and increased angiogenic growth factor secretion in both conditions. The regenerative potential of the miRNA-i-activated scaffold *via* macrophage polarisation was validated through inflammatory and angiogenic functional assays with endothelial cells. In parallel, scaffold-mediated miRNA-155 delivery to dorsal root ganglia (DRG) promoted neurogenic outcomes through enhanced axonal regrowth, essential for the synergistic repair of chronic wounds across the skin-nerve axis. *In vivo* implantation of miRNA-155 inhibitor-activated scaffolds in chicks demonstrated successful integration without disrupting vascular network formation. Collectively, these findings establish the miRNA-155 inhibitor-activated scaffold as a regenerative platform with anti-inflammatory, pro-angiogenic, and neurogenic outcomes, offering a multifaceted solution for chronic wound healing applications.

**Graphical Abstract:** 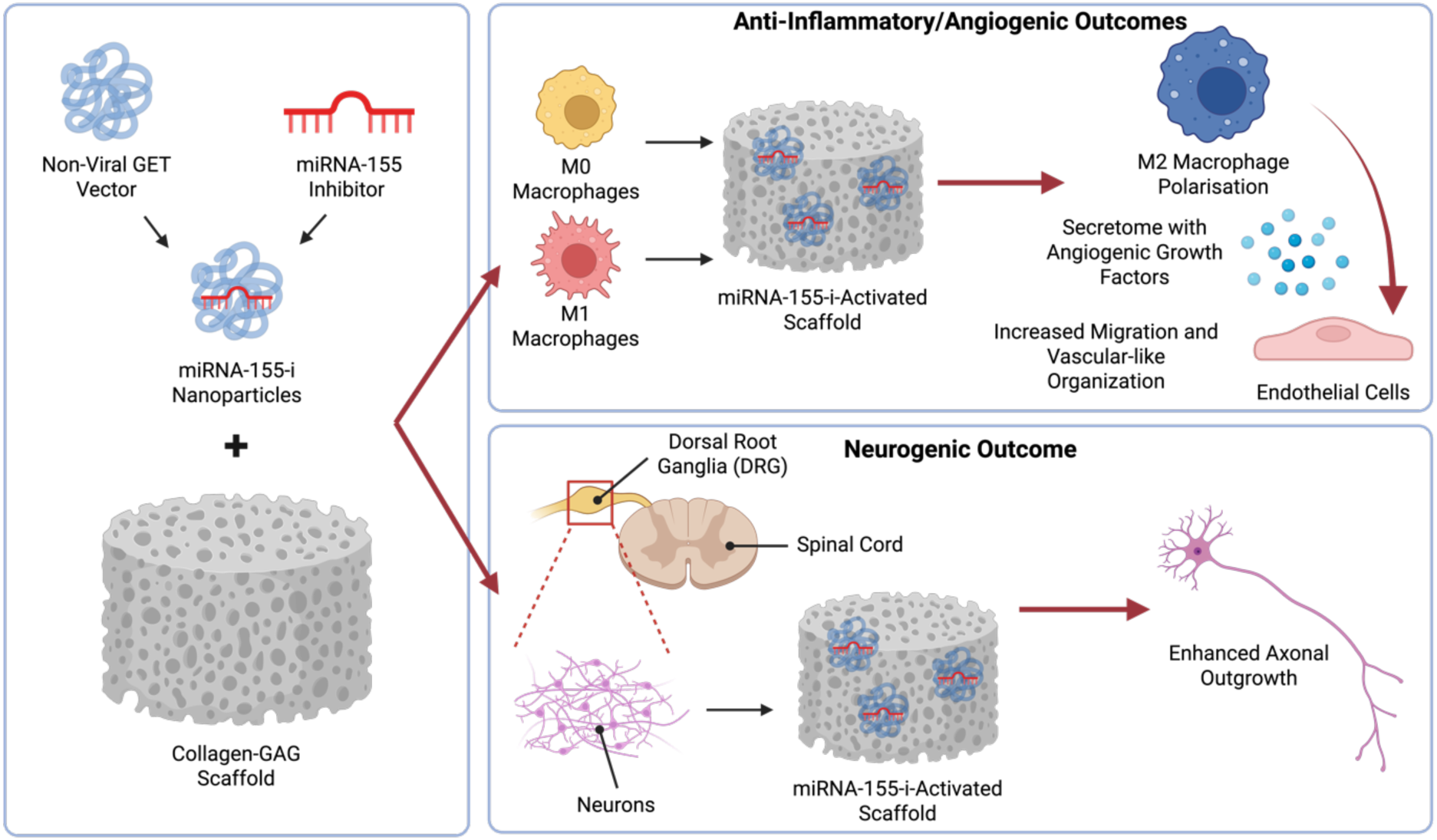

## 1. Introduction

Chronic wounds, including diabetic foot ulcers, represent a significant clinical challenge due to their persistent inflammatory microenvironment and impaired progression through the normal healing stages.[1], [2] Across Europe, an estimated two million individuals are affected by chronic wounds – a figure projected to increase with ageing demographics and the prevalence of obesity and diabetes.[3] In Ireland, chronic wounds impact approximately 1.5% of the population, contributing to an annual economic burden of €630 million EUR,[4] underscoring the need for therapeutic strategies capable of restoring the healing process.

Under healthy physiological conditions, skin injury initiates the wound healing cascade consisting of four overlapping and sequential stages, namely haemostasis, inflammation, proliferation, and remodelling.[5] This coordinate response ensures the restoration of the structural and functional integrity of skin. However, underlying co-morbidities, such as diabetes mellitus, can dysregulate this process leading to the formation of chronic non-healing wounds.[6] These injuries present a dysregulated inflammatory microenvironment driven by excessive immune cell infiltration and sustained pro-inflammatory signalling, which exacerbates the pathological state.[7], [8] Moreover, compromised nerve regeneration due to inflammation reduces beneficial cell interactions across the skin-nerve axis.[9], [10], [11], [12]

Macrophages are prominent among these recruited immune cells as highly plastic cells capable of transitioning between pro-inflammatory (M1) and anti-inflammatory (M2) phenotypes in response to environmental signals.[13] In physiological wound healing, M1 macrophages initially mediate cell debris and bacteria removal.[14] This phase is then followed by a phenotypic switch towards a M2 macrophage phenotype that supports tissue repair through the secretion of regenerative growth factors, promotion of extracellular matrix (ECM) deposition, stimulation of angiogenesis, and nerve regeneration.[14], [15] However, the M1 phenotype persists in chronic wounds, preventing the resolution of inflammation and subsequent healing, thereby leaving wounds open and susceptible to infections.[16], [17], [18]

Standard therapeutic strategies used for the treatment of chronic wounds include wound debridement, antibiotics, and the implantation of passive dermal substitutes, such as the Integra dermal regeneration template (Integra® DRT) which show varying potential to facilitate the healing process.[19] Collectively, these interventions have shown limited efficacy and inconsistent outcomes across patients, particularly in complex and recurring wounds.[1], [3], [20], [21], [22] Critically, current treatments do not directly target the disrupted inflammatory response nor do they address coexistent – yet often overlooked – challenges such as impaired nerve regeneration.[23]

Combinatorial therapeutic strategies utilising biomaterial scaffolds and gene delivery to enhance regenerative responses have shown potential to offer a multifaceted approach to wound healing.[24], [25], [26], [27], [28] In particular, collagen-glycosaminoglycan (CG) scaffolds have shown great potential to support wound closure while allowing the incorporation of different DNA-or RNA-based therapeutics that modulate genes of interest, thereby influencing cellular behaviour and enabling healing.[24] Among these nucleic acids, non-coding microRNAs (miRNAs) are of interest due to their role in regulating multiple genetic targets through post-transcriptional RNA interference.[29], [30] Crucially, miRNAs are highly expressed in every stage of wound healing and influence pathological processes when dysregulated, including persistent inflammation, M1 macrophage overactivation, decrease vascularisation, and impaired nerve regeneration observed in chronic wounds.[31], [32]

Among the multiple miRNAs involved in inflammation, miRNA-155 stands out as a well-characterised, multifunctional miRNA with essential roles in macrophage activation and phenotypic polarisation.[33], [34], [35] MiRNA-155 expression is strongly up-regulated through nuclear factor-kappa-light-chain-enhancer of activated B cells (NF-κB) signalling as a response to inflammatory stimuli such as lipopolysaccharide (LPS) and interferon-gamma (IFN-γ).[35] Functionally, miRNA-155 represses the suppressor of cytokine signalling 1 (SOCS1), Src homology 2 domain containing inositol polyphosphate 5-phosphatase (SHIP1), and B-cell lymphoma 6 (BCL6) which, in turn, accentuate inflammatory pathways.[36], [37], [38] Under pathological conditions, miRNA-155 expression is persistently up-regulated contributing to M1 macrophage maintenance and exacerbation of the chronic state.[35]

Importantly, the effects of miRNA-155 modulation extend beyond its well-established role in inflammation. Evidence from several *in vitro* and *in vivo* studies indicate that neurons in the central and peripheral nervous systems regain regenerative potential following miRNA-155 inhibition.[39], [40], [41], [42] For example, in the peripheral nervous system, miRNA-155 knockdown has been shown to promote spontaneous axon growth and reduce neuronal toxicity,[39], [40] highlighting the multi-faceted capability of miRNA and the beneficial outcome it can provide through healing of chronic wounds across the skin-nerve axis.

Given the multifaceted role of miRNA-155 in wound healing, inhibition of this miRNA represents a key therapeutic target for the resolution of the chronic wound environment. However, the clinical translation of gene delivery still faces challenges – most notably, the poor cellular uptake and rapid degradation of free nucleic acids under physiological conditions.[43], [44] Thus, the selection of an appropriate carrier vector is crucial for the protection of the genetic material, targeting efficiency, and overall safety of the delivery system. Although viral vectors offer superior efficiency, non-viral vectors have shown lower immunogenic risks and great scalability potential.[5], [44], [45] In particular, the non-viral glycosaminoglycan enhanced transduction (GET) peptide has proven great cellular uptake by integrating cell penetrating and heparan sulphate GAG-binding sequences, making it a suitable carrier for miRNA inhibitor delivery.[46], [47], [48], [49], [50]

The objective of this study involved the development of a miRNA-155 inhibitor-activated scaffold (CG-155-i) for localised gene delivery, leading to regenerative macrophage polarisation, enhanced vascularisation, and the regeneration of injured peripheral neuron axons in chronic wound healing. Initially, this study focused on the formulation of the GET encapsulated miRNA-155 inhibitor delivery system and incorporation with the CG scaffold platform. Thereafter, the ability of the CG-155-i framework to induce anti-inflammatory macrophage polarisation in non-polarised and pro-inflammatory immune cells was assessed. This was followed by the profiling of CG-155-i-mediated macrophage cytokine release and its influence on inflammatory and angiogenic processes in endothelial cells. Additionally, the pro-neurogenic ability of the platform to positively influence axon regrowth was characterised in an *ex vivo* dorsal root ganglia model of axonal injury. Finally, the *in vivo* biocompatibility of the CG-155-i platform was evaluated in a chicken chorioallantoic membrane (CAM) model.

## 2. Materials and Methods

All reagents were purchased from Thermo Fisher Scientific (Ireland) unless otherwise stated. All cell culture was performed at 37 °C and 5% CO_2_ unless otherwise stated.

### 2.1. Development of miRNA inhibitor nanoparticles

The miRIDIAN microRNA hairpin inhibitor hsa-miRNA-155-5p (miRNA-155 inhibitor) and scramble hairpin inhibitor negative control (miRNA-Scr inhibitor) (Dharmacon, UK) were combined with the glycosaminoglycan enhanced transduction (GET) peptide through electrostatic interactions to form complexes at charge ratio 6 (CR6) as previously established.[24] miRIDIAN microRNA hairpin inhibitor red transfection control (Dharmacon, UK) was complexed in the same manner as above to enable fluorescent tracking of nanoparticles (Cy3 NPs).

### 2.2. Physicochemical characterisation of miRNA inhibitor nanoparticles

Physicochemical characterization of the miRNA-155 inhibitor nanoparticles was carried out by dynamic light scattering (DLS) (Zetasizer 3000 HS, Malvern, UK) and nanoparticle tracking analysis (NTA) (NanoSight NS300, Malvern, UK) to determine charge (zeta potential), polydispersity index (PDI), and size distribution, respectively. For the analysis of zeta potential and PDI, nanoparticles were prepared with molecular grade water (MG H_2_O), the volume was increased to 1 mL and transferred to a disposable folded capillary cell (Malvern, UK) before analysis. For size distribution measurements, nanoparticles were also prepared as before, and data was captured with a sCMOS camera and Blue488 laser. Data evaluation was carried with NTA 2.3 software (Malvern, UK).

Complexation efficiency was measured by preparing the miRNA-155 inhibitor nanoparticles (40 pmol miRNA inhibitor) as previously mentioned. Then, free miRNA concentrations were measured using a Quant-it™ RiboGreen Reagent and RNA Assay kit following the manufacturer’s protocol. RNA concentrations were extrapolated from the standard curve and the percentage of not complexed miRNA was determined by normalising the measurements against a free miRNA control (40 pmol miRNA inhibitor).

### 2.3. Assessment of miRNA-i nanoparticle transfection efficiency

To characterise the transfection efficiency of miRNA-I nanoparticles on macrophages, human THP-1 monocytic cells were culture with varying concentrations of Cy3-tagged miRNA-i nanoparticles over 3 days. Initially, 1.5 x10^4^ THP-1 cells were seeded on 13 mm coverslips (Cat# 17274914, Fisher Scientific, UK) within 12-well culture plates and conditioned overnight with phorbol 12-myristate 13-acetate (PMA, 20 ng mL^-1^) in Roswell Park Memorial Institute (RPMI) 1640 culture medium containing 10% FBS and 1% P/S. Pro-inflammatory (M1) macrophage differentiation was achieved by incubating the cells with 1 mL of RPMI medium supplemented with interferon-γ (IFN-γ, 5 ng mL^-1^) and liposaccharide (LPS, 100 ng mL^-1^, Merck, Ireland) for 72 hours.

After 24 hours from PMA induction, varying concentrations of Cy3-tagged miRNA-i nanoparticles and free Cy3-tagged miRNA-i were delivered to the cells and cultured for 72 hours. Following this, cells were fixed in 4% PFA for 1 hour at 4°C before being washed 3 times with DPBS and stored at 4°C until further processed. Cells were then permeabilised with 0.1% Triton X-100 solution for 5 mins followed by incubation with Alexa Fluor 488 Phalloidin (1:500, Cat # 49409, Merck, Ireland) for 1 h. Finally, cells were incubated with Hoechst 33342 (1:10000) for 15 min with three DPBS washes in between steps. Coverslips were then mounted in glass slide with Fluoromount-G™ Mounting Medium (Cat# 00-1958-02). Coverslips were imaged using a Nikon Eclipse 90i microscope, maintaining consistent gain, exposure, and magnification. Images were analysed using FIJI software[51] to calculate nuclei count, cell coverage, Cy3 expression, and transfected cells using automated scripts.

### 2.4. Collagen-GAG (CG) scaffold fabrication, crosslinking, and functionalisation with miRNA inhibitor nanoparticles

The CG slurry used for scaffold fabrication was prepared as previously described.[52] After lyophilisation, CG scaffolds were chemically crosslinked with 1-ethyl-3-(3-dimethyl aminopropyl)-carbodiimide (EDAC) and N-hydroxy succinimide (NHS) (Merck, Germany) to enhance their structural properties following a previously established method.[24], [52] After crosslinking, scaffolds were washed 3 times with DPBS to remove excess EDAC/NHS before sterilization in 70% ethanol for 30 min. Finally, scaffolds were washed 3 more times with DPBS under sterile conditions. CG scaffolds were then gene-activated through the soak-loading of miRNA-155 inhibitor (CG-155-i) or miRNA-Scr inhibitor (CG-Scr-i) nanoparticles. Briefly, nanoparticles were soak-loaded on one side of the CG scaffolds (20 pmol miRNA inhibitor) and incubated for 45 min at 37°C. Then, scaffolds were flipped, and the process was repeated.

### 2.5. Morphology, distribution, and release profile of miRNA-i-activated scaffolds

To confirm that functionalisation of CG scaffolds with miRNA nanoparticles did not affect the microstructure, scaffolds were imaged using a scanning electron microscope (SEM) as previously described.[53] Briefly, CG and CG-155-i scaffolds were gene-activated and lyophilised through a critical pressure sublimation step. Dried scaffolds were bisected using a scalpel, followed by mounting on metallic pin studs with carbon tape. Mounted scaffolds were sputtered with a 80/20 mixture of gold/palladium alloy to a thickness of ∼4 nm in a Cressington 108 auto sputter coater. Scaffold microstructure was assessed using a Zeiss Ultra FE-SEM (Zeiss, Germany) with an accelerating voltage of 3kV at several magnifications.

To further confirm the distribution of nanoparticles on the scaffolds, fluorescently tagged miRNA (Cy3) NPs were soak-loaded on CG (CG-Cy3) scaffolds as mentioned above. Cross-sections of these scaffolds were then visualised using a Zeiss LSM 710 confocal microscope.

To assess the release profiles from miRNA-i-activated scaffolds, miRNA-155 inhibitor nanoparticles were formulated in MG H_2_O before soak loading on scaffolds (0.2 nmol). Samples were then placed in 24-well plates and flooded with 2 mL MG H_2_O. The release assessment was carried out in static conditions at 37 °C. At every timepoint, 200 μL of supernatant were collected and replaced with 200 μL of fresh MG H_2_O. The supernatant containing released nanoparticles was then incubated with 50 μL of heparin (1 mg mL^-1^) for 90 min to promote the dissociation of nanoparticles. Finally, the amount of miRNA released was quantified using a Quant-it™ RiboGreen Reagent and RNA Assay kit following the manufacturer’s protocol.

### 2.6. Cell culture, differentiation, polarisation, and seeding on miRNA-i-activated scaffolds

Normal human dermal fibroblasts (HDFs) from adult donors were purchased from PromoCell (Germany) and cultured in growth media containing low glucose (1.0 g L^-1^) Dulbecco’s Modified Eagles Medium (DMEM) supplemented with 10% FBS and 1% penicillin-streptomycin (P/S).

Human umbilical vein endothelial cells (HUVECs) were purchased from Lonza (Switzerland) and cultured in endothelial growth medium-2 (EGM-2) supplemented with SupplementMix (PromoCell, Germany).

Human THP-1 monocytic cells (ATCC – TIB-202, USA) were cultured in Roswell Park Memorial Institute (RPMI) 1640 culture medium containing 10% FBS and 1% P/S. To induce non-polarised (M0) macrophage differentiation on scaffolds, 1.25 x 10^5^ cells were seeded onto the first side of the gene-activated scaffolds at a concentration of 2.5 x 10^6^ cells mL^-1^ in the presence of phorbol 12-myristate 13-acetate (PMA, 20 ng mL^-1^). Scaffolds were then allowed to incubate at 37°C for 15 mins before repeating the process on the opposite side. Finally, all the wells were flooded with 1 mL of PMA-containing RPMI medium for 18 hours. Pro-inflammatory (M1) macrophage differentiation was achieved by following the same steps used for M0 differentiation with the additional incubation of macrophage-seeded scaffolds in 1 mL of RPMI medium supplemented with interferon-γ (IFN-γ, 5 ng mL^-1^) and liposaccharide (LPS, 100 ng mL^-1^, Merck, Ireland) for 72 hours.

### 2.7. Assessment of cell viability post-transfection on miRNA-i-activated scaffolds over 7 days

To understand the effect of scaffold-mediated miRNA-155 inhibitor transfection on HDFs, HUVECs, and THP-1 macrophages viability, cell-based assays were carried out. Following cell transfection on scaffolds, cell metabolic activity was determined through an Alamar Blue™ Cell Viability assay according to the manufacturer’s protocol. Briefly, media was removed and growth medium (DMEM, EGM-2, or RPMI, respectively) containing 10% Alamar Blue™ reagent was added to the cells (1 mL). Samples were then incubated for 2 h at 37°C in the dark. The supernatant was collected, and the fluorescence of each sample was measured in triplicate (ex: 570 nm, em: 585 nm) using an Infinite® 200 PRO plate reader (Tecan Group Ltd., Switzerland). Fluorescence measurements of the miRNA inhibitor transfected groups were normalised to the miRNA-free (CG) control.

DNA content was measured with a Quant-iT™ PicoGreen™ dsDNA Assay Kit. Media was removed and wells were flooded with 1 mL buffer (0.2 M sodium carbonate + 0.1% Triton X-100 in DI H_2_O) to lyse the cells. Samples were then subjected to 3 freeze-thawing cycles at -80 °C before measurements were carried out. Fluorescence measurements (ex: 480 nm, em: 520 nm) were performed using an Infinite® 200 PRO plate reader. Finally, the DNA concentration was extrapolated from the standard curve.

### 2.8. Enzyme-linked immunosorbent assay (ELISA) for growth factor quantification post-transfection

A human vascular endothelial growth factor (VEGF) ELISA kit (R&D Systems, USA) was used to quantify the protein release from cells transfected on gene-activated scaffolds. ELISAs were carried out as previously reported[54] with the conditioned media collected on days 1, 3 and 7 post-transfection. Absorbance measurements at 450 and 540 nm were taken using an Infinite® 200 PRO plate reader. Finally, VEGF expression was calculated by extrapolation from the standard curve.

### 2.9. Gene expression analysis using qRT-PCR

RNA expression was assessed through quantitative real time PCR (qRT-PCR). After 3 days of culture on scaffolds, RNA was extracted from macrophages by lysing cells using 500 μL QIAzol lysis reagent (Qiagen, Ireland) per scaffold, and subsequently extracting the RNA fraction using a miRNeasy kit (Qiagen, Ireland) according to the manufacturer’s protocol. cDNA templates were then produced for RNA and microRNA analysis using a QuantiTect Reverse Trancription Kit (Qiagen, Ireland) and a Taqman™ Advanced miRNA cDNA synthesis kit, respectively, following the manufacturer’s protocol. qRT-PCR was carried out with a variety of target genes (Supp. Table 1) using a Lightcycler 480 II (Roche, UK). Finally, the ΔΔCt method was utilised to calculate the fold change expression of the genes of interest. The sustained inhibition of miRNA-155 was also validated after 7 days under M0 and M1 conditions following the RT-PCR procedure for microRNA analysis.

### 2.10. Immunostaining, imaging, and morphological analysis of cell-seeded miRNA-i-activated scaffolds

Cell morphology, distribution, and marker expression of THP-1 macrophages, HDFs, and HUVECs on miRNA-i-activated scaffolds were analysed using mouse anti-CD206 antibody (Cat# sc-376108 AF488, Santa Cruz Biotechnologies, USA), mouse anti-CD80 antibody (Cat# MA5-15512), mouse anti-CD144 antibody (Cat# 14-1449-82), F-actin cytoskeleton, and cell nuclei visualisation. Briefly, after 3 or 7 days of culture, all scaffolds were fixed in 4% PFA for 1 hour at 4°C before being washed 3 times with DPBS and stored at 4°C until further processed. For immunostaining, the scaffolds were permeabilised with 0.1% Triton X-100 solution for 5 mins followed by blocking at room temperature (RT) with 1% bovine serum albumin (BSA) in DPBS for 2 hours. Following blocking, macrophage-seeded scaffolds were incubated with mouse anti-CD206 antibody (1:200) or mouse anti-CD80 antibody (1:200) in 1% BSA overnight at 4°C. The next day, scaffolds were washed and incubated with Alexa Fluor 555 Phalloidin™ (1:500) and goat anti-mouse Alexa Fluor 488™ secondary antibody (1:1000) for 2 hours and Hoechst 33342 (1:10000) for 15 min with three DPBS washes in between steps. HDF-seeded scaffolds were fixed and permeabilised as before followed by incubations with Alexa Fluor 555 Phalloidin™ and Hoechst 33342. HUVEC-seeded scaffolds were also fixed, permeabilised, and blocked before incubation with mouse anti-CD144 (1:200) overnight at 4°C, followed by Alexa Fluor 555 Phalloidin™ and Hoechst 33342 staining. All scaffolds were imaged using a Zeiss LSM 710 confocal microscope, maintaining consistent gain, exposure, and magnification. Images were analysed using FIJI software[51] to calculate nuclei count, cell coverage, circularity and expression of CD206 (from M0 macrophages), CD80 (from M1 macrophages), and CD144 (HUVECs) using automated scripts.

### 2.11. Assessment of paracrine signalling from miRNA-i-activated scaffolds on inflammatory and angiogenic processes

To characterise the expression of angiogenic and inflammatory cytokines released by conditioned macrophages, a cytokine array analysis was performed with supernatant collected from M0 and M1 macrophage-seeded scaffolds on day 7 post-transfection. The assay was carried out using a 43-target Human Angiogenesis Antibody Array Membrane (ab193655, Abcam, UK) according to the manufacturer’s protocol. Densitometry measurements were then obtained in FIJI, and each cytokine expression was normalised using the M0 CG group as a control.

#### Assessment of anti-inflammatory effect from miRNA-i-activated scaffolds

To analyse the effect of scaffold-driven macrophage polarisation on the expression of pro-inflammatory marker intercellular cell adhesion molecules (ICAM) in endothelial cells, a functional assay was devised through the Transwell co-culture of THP-1 macrophages and HUVECs. Initially, 6.0 x 10^4^ HUVECs were seeded on 13 mm round coverslips (Cat# 17274914, Fisher Scientific, UK) in 12-well plates and conditioned with LPS (40 ng mL^-1^) in EGM-2 (1 mL) for 24 hours. In parallel, THP-1 cells were-seeded and differentiated towards M0 macrophages on miRNA-i-activated scaffolds as outlined above. The next day, media was removed from the monolayers and scaffolds, Transwell inserts were placed on top of the monolayer with THP-1 seeded scaffolds sitting on the membrane. Finally, M0 media (RPMI) or M1 media (IFN-γ/LPS-conditioned RPMI) was added to the Transwell systems (1 mL).

After 3 days of culture, the Transwell inserts and scaffolds were removed and HUVEC monolayers were fixed in 4% PFA for 15 min at RT. Monolayers were permeabilized with 0.1% Triton X-100 solution for 5 mins followed by blocking at RT with 1% BSA in DPBS for 2 h. Following blocking, HUVECs were incubated with mouse anti-ICAM-1 antibody (1:200, Cat# MA5407) in 1% BSA overnight at 4°C. The next day, the monolayers were incubated with Alexa Fluor 555 Phalloidin™ (1:500) and Alexa Fluor 488™ secondary antibody (1:1000) for 1 h and Hoechst 33342 (1:10000) for 15 min with 3 DPBS washes in between steps. Finally, HUVECs on coverslips were mounted on glass slides with Fluoromount-G™ Mounting Medium (Cat# 00-1958-02) and imaged with a Nikon Eclipse 90i microscope maintaining consistent gain, exposure, and magnification. Images were then analysed in FIJI to determine nuclei count, cell coverage, and ICAM-1 expression.

#### Assessment of pro-angiogenic effect from miRNA-i-activated scaffolds

To assess the capability of miRNA-i-activated scaffolds to enhance migration and organization of endothelial cells through paracrine signalling, scratch and tube formation assays were performed with the conditioned media of macrophage-seeded scaffolds 7 days post-transfection. The effect on endothelial cell migration was assessed through a scratch assay using a previously established protocol.[24] Controls including endothelial basal medium (VEGF-), RPMI with 10% FBS supplementation (FBS+) and endothelial growth medium (VEGF+) were also used before transferring the plate to a Zeiss Celldiscoverer 7 microscope. Images were taken every hour for 48 h. Cell migration was calculated as the change in scratch area relative to the start timepoint (0 h).

To characterise the effect of miRNA-i-activated scaffolds on endothelial cell organisation, a tube formation assay was carried out as previously described.[24] Briefly, Geltrex™ LDEV-Free Reduced Growth Factor Basement Membrane Matrix was thawed 24 h prior to use at 4°C. Then, each well of a 96-well cell culture plate was coated with 80 μL of Geltrex on each well and centrifuged before gelation at 37°C for 30 mins. Next, 6.0 x 10^4^ HUVECs were resuspended in 600 μL of conditioned media from each group and controls. After incubation, 200 μL of conditioned media cell-suspensions were added to the Geltrex-coated wells in triplicate. The plate was transferred to a Zeiss Celldiscoverer 7 microscope and images were taken every hour for 48 h. Image analysis was carried out using the Angiogenesis Analyzer for ImageJ[55] to determine the junction number, branch number, isolated segments, and tube length.

### 2.12. Characterisation of pro-neurogenic ability of miRNA-i-activated scaffolds in *ex vivo* axonal injury model

To assess the ability of the miRNA-i-activated scaffolds to support axonal growth in a suitable *ex vivo* model of axon injury, dorsal root ganglia (DRGs) from 5-month-old adult female C57BL6 mice were seeded onto the scaffolds following a previously established method.[24] DRG post-mortem harvesting was carried out under HPRA individual license (AE19127/I259) and with ethical approval from the RCSI research Ethics Committee (REC202211008). For each DRG, the roots were first trimmed of its associated nerves using fine micro-scissors and then placed individually in the middle of each scaffold and immersed in 400 μL of Neurobasal™ medium supplemented with 1% P/S, 1% Glutamax, and 2% B27 supplement (DRG culture medium) and incubated for 12 h to allow the DRGs to adhere. Following this, the volume of media was increased to 750 μL to fully cover the scaffold and DRG and the media (50%) was replaced every second day. On days 3, 7, and 14, media was collected before collecting the scaffolds on day 14. For fixation, scaffolds were washed twice with DPBS and fixed in 4% PFA for 1 h at RT before being washed and incubated overnight at 4°C in an anti-mouse β-tubulin III antibody (1:500) in DPBS containing 0.% Triton X-100. Finally, nuclei were stained with Hoechst 33342 (1:10000) for 15 min with three DPBS washes in between steps. The scaffolds were imaged using a Zeiss LSM 710 confocal microscope under consistent gain, exposure and magnification. Analysis of neurite length was carried out by calculating the straight-line distance from axonal growth cone to the DRG body using the manual tracing tool function in FIJI.

Quantification of lactate dehydrogenase (LDH) release from DRGs was carried out with the conditioned media collected on days 3, 7, and 14 from the DRGs on miRNA-i-activated scaffolds. Following collection, media was frozen at -80°C and thawed just before performing the assay with a CyQuant™ LDH Cytotoxicity Assay (Cat# C20300) according to the manufacturer’s protocol.

### 2.13. Assessment of miRNA-i-activated scaffolds in an *in vivo* chicken chorioallantoic membrane model

To assess the biological response from the miRNA-i-activated scaffolds in an *in vivo* model, a chicken chorioallantoic membrane (CAM) assay was performed. Initially, fertilised chicken eggs (Shannon Vale Foods, Ireland) were cleaned and incubated at 37°C in a humid environment for 3 days. Following this, eggshells were cracked, and the contents were transferred to 100 mm Falcon™ Standard Tissue Culture Dishes (Cat# 10212951, Fisher Scientific, UK) and incubated for a further 4 days. Then, three scaffolds of each group were implanted on the membrane by gently placing each scaffold on one of the larger vessels extending from the chicken embryo. Subsequently, embryos were incubated for a further 5 days before imaging the formed vasculature with a Nikon D5600 camera connected with a Nikon AF-P DX Nikkor 18-55 mm f/3.5-5.6G lens at a fixed distance and magnification. Finally, scaffolds were gently removed from the membrane and fixed in 4% PFA overnight at 4°C before histological analysis and the chicken embryos were immediately euthanised by decapitation and clipping of central vasculature.

Analysis of the change in vascularisation was carried out on membrane areas surrounding scaffolds in the images obtained from chicken embryos with an automated script in Fiji software. Quantification of vessel length, branch number, and junction number was compared among scaffold groups and scaffold-free embryos (NT).

Following collection and fixation of scaffolds, samples were processed overnight in an automatic tissue processor (ASP300, Leica, Germany), dissected, and embedded in paraffin wax blocks to visualise membrane integration within the scaffolds. Thereafter, the blocks were sectioned using a rotary microtome (Microsystems GmbH, Germany) into 10 μm sections and mounted on poly-L-lysine-coated glass slides. Wax in sections was melted overnight in an oven at 60°C before rehydration in a graded series of ethanol and staining with Haematoxylin & Eosin (Cat# HHS32-1L and HT110232-1L, H&E, Merck, Ireland) and Masson’s Goldner trichrome staining kits (Cat# 1004850001, Merck, Ireland) following the protocols presented in the Supplementary Methods. Finally, the stained slides were finally mounted overnight with DPX mountant for histology (Cat# 06522, Merck, Ireland), allowed to dry overnight at RT before being imaged using Nikon Eclipse 90i microscope with consistent magnification and exposure. Final processing was carried out in Fiji software.

### 2.14. Statistical Analysis

Statistical analysis was carried out using Graph-Pad Prism v 10.4.1. One-way ANOVA with a Tukey post-hoc test was used when more than one treatment was compared. Two-way ANOVA with Bonferroni post-hoc test was used when more than one treatment was compared across two factors. All experiments were performed at least in triplicate with three replicates or more. Results are expressed as mean ± standard deviation (SD).

## 3. Results

### 3.1. miRNA-155-i nanoparticles present suitable properties for cellular internalisation on the collagen-GAG scaffolds

The physicochemical properties of nanoparticles are key features that have a high impact on cellular uptake and internalisation processes such as endosomal escape or lysosomal degradation.[56] Therefore, assessment of miRNA-155 inhibitor/GET (miRNA-155-i) nanoparticles was carried out through analysis of size distribution, charge, polydispersity index (PDI), and complexation efficiency, revealing a suitable diameter and charge for cellular transfection. MiRNA-155 inhibitor nanoparticles exhibited an average diameter of 86.9 ± 35.4 nm (Fig. 1A) and a positive charge of 27.5 ± 1.9 mV (Fig. 1B). Additionally, the PDI of the miRNA-155 inhibitor nanoparticles was measured as 0.28 ± 0.06 with a complexation efficiency of 98.1 ± 0.7 %.

**Figure 1.**
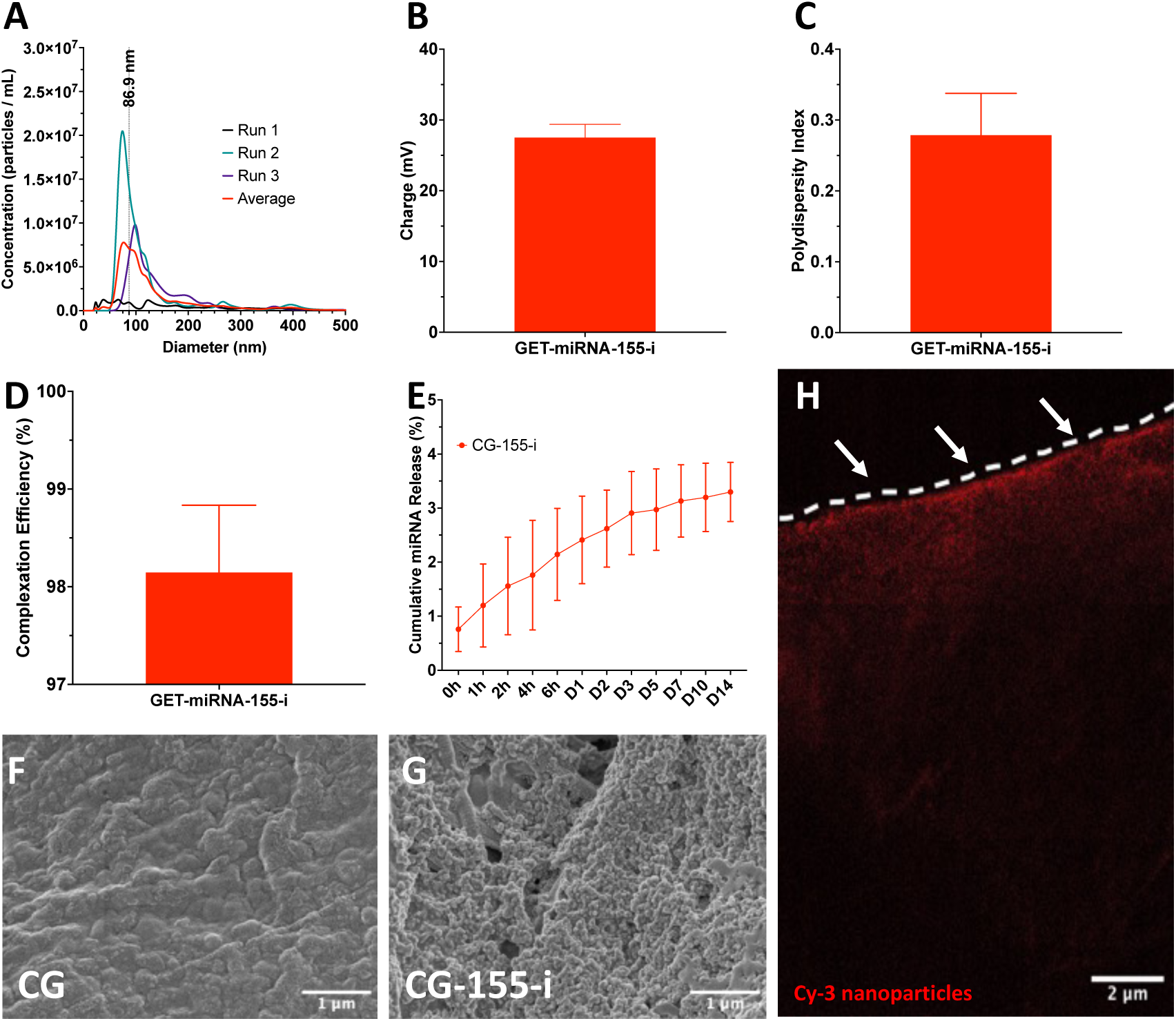
MiRNA-155 inhibitor nanoparticles present optimal physicochemical properties and homogeneous distribution on the surface of collagen-GAG (CG) scaffolds. A-C) MiRNA-155 inhibitor nanoparticles exhibited an average diameter of 86 nm, a positive charge of 27 mV, and a PDI of 0.28. D) Complexation efficiency measurements revealed a high degree of complexation of 98.1%. E) The nanoparticle release profile obtained from CG-155-i scaffolds shows a gradual cumulative release of nanoparticles up to 14 days. F-G) Scanning Electron Microscope (SEM) images show that nanoparticles retain a spherical morphology on the porous microstructure of the scaffolds. H) Confocal imaging of Cy-3-tagged miRNA nanoparticles displays an even nanoparticle distribution on the CG scaffolds’ surface – segmented line represents scaffold surface. Data shows mean ± SD (n=3).

Subsequent characterisation of the release profile from collagen-GAG (CG) scaffolds soak-loaded with miRNA-155 inhibitor nanoparticles (CG-155-i) revealed a gradual release of cargo with that more than 95% retained within the porous structure for up to 14 days (Fig. 1E) while evaluation of miRNA-155 inhibition revealed a sustained downregulation of the target miRNA for up to 7 days (Supp. Fig. 1). Additionally, assessment of transfection efficiency displayed a dose-dependent increase in transfection in macrophages of up to ∼60 % (Supp. Fig. 2). Evaluation of scaffold porosity through scanning electron microscope (SEM) imaging showed that the porous architecture of the CG scaffolds was not affected after nanoparticle soak-loading while miRNA-155 inhibitor nanoparticles maintained a spherical morphology within the pores (Fig. 1F-G). Moreover, confocal visualisation of nanoparticle distribution within CG scaffold cross-sections with Cy3-tagged miRNA revealed a homogeneous nanoparticle distribution on the scaffolds’ surface (Fig. 1H).

### 3.2. CG-155-i scaffolds drive a regenerative phenotype from non-polarised macrophages

Having established the suitable physicochemical nanoparticle properties and their distribution on CG scaffolds, the effect of the miRNA-i-activated scaffolds on dermal cells was initially investigated. Human dermal fibroblasts (Supp. Fig. 3) and endothelial cells (Supp. Fig. 4) were seeded on miRNA-i-activated scaffolds and no detrimental effects on metabolic activity, DNA content, or cell morphology were observed over 7 days of culture on the CG-155-i scaffolds. However, downregulation of VE-cadherin was observed on endothelial cells when seeded on CG-Scr-i scaffolds (Supp. Fig. 4).

Next, the influence of miRNA-i activation on macrophage polarisation was assessed by seeding monocyte-derived non-polarised (M0) macrophages on the scaffolds. Scaffold-mediated miRNA-155 inhibition led to an increase in metabolic activity (Fig. 2A) and DNA content (Fig. 2B) for up to 7 days compared to M0 cells grown on the miRNA-free CG and miRNA scramble inhibitor-activated (CG-Scr-i) scaffolds. Moreover, M0 macrophages on CG-155-i scaffolds produced a significantly higher release of the pro-angiogenic growth factor vascular endothelial growth factor (VEGF) than the macrophages grown on miRNA-free CG scaffolds on days 3 and 7 post-transfection (Fig. 2C). However, VEGF expression on the CG-Scr-i group matched that of the CG-155-i scaffolds on day 7.

**Figure 2.**
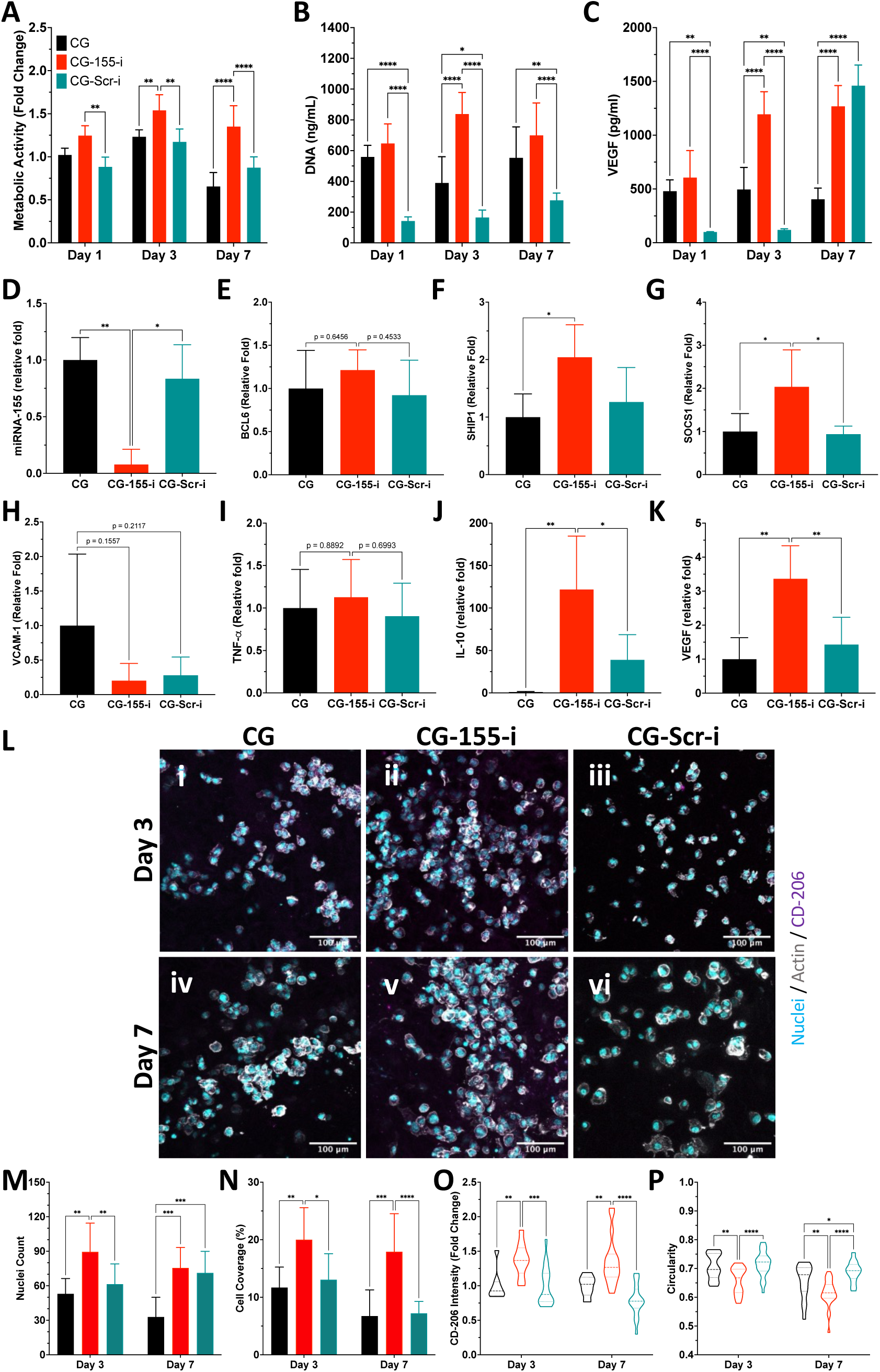
Non-polarised (M0) macrophages grown in CG-155-i scaffolds are driven towards an anti-inflammatory (M2) phenotype. A-B) Assessment of cell viability using metabolic activity and DNA content showed increased macrophage activity and proliferation on the CG-155-i group over 7 days. C) Quantification of VEGF release was significantly enhanced in macrophages seeded on CG-155-i scaffolds. D-G) Gene expression analysis of miRNA-155 and directly regulated target genes SHIP1, BCL6, and SOCS1 demonstrate the activation of anti-inflammatory processes following miRNA-155 inhibition via SHIP1 and SOCS1. H-K) Gene quantification of inflammatory and angiogenic-associated genes VCAM-1, TNF-α, IL-10, and VEGF showed anti-inflammatory/pro-angiogenic response from M0 macrophages on CG-155-i scaffolds. L-P) Confocal visualisation of M0 macrophages on scaffolds exhibited higher macrophage count and coverage with increased CD-206 intensity and circularity on CG-155-i scaffolds. Data shows mean ± SD (n=5), * indicates p<0.05, ** p<0.01, *** p<0.001, **** p<0.0001.

Following cytokine expression and cell viability characterisation, miRNA-155 expression and its effect on downstream and inflammatory genes were quantified on day 3 post-transfection. Initially, miRNA-155 inhibition was confirmed through the significant downregulation of the target gene in the CG-155-i group compared to the macrophages on CG and CG-Scr-i scaffolds (Fig. 2D). Subsequent quantification of the miRNA-155 downstream target B-cell lymphoma (BCL6, Fig. 2E) did not show a significant change on the CG-155-i group compared to CG and CG-Scr-i scaffolds. However, measurement of the downstream targets Src homology 2 domains containing inositol polyphosphatase 5-phosphatase 1 (SHIP1, Fig. 2F) and suppressor of cytokine signalling (SOCS1, Fig. 2G) displayed significantly increased expressions of ∼2 fold compared to CG scaffolds, suggesting an anti-inflammatory downstream effect on macrophages following miRNA-155 inhibition. Further expression analysis of genes involved in inflammation, including vascular cell adhesion molecule (VCAM-1, Fig. 2H) and tumour necrosis factor-alpha (TNF-α, Fig. 2I) showed a trending decrease. In contrast, interleukin-10 (IL-10, Fig. 2J) expression was increased more than 100-fold in macrophages grown in CG-155-i scaffolds compared to both CG and CG-Scr-i. Moreover, expression of the pro-angiogenic gene VEGF was significantly increased ∼3 fold in the CG-155-i group (Fig. 2K), highlighting the potential anti-inflammatory/pro-angiogenic effect of miRNA-155 inhibition in M0 macrophages through CG-155-i scaffolds.

Subsequent visualisation and analysis of M0 macrophages on miRNA-i-activated scaffolds revealed an increased expression of the anti-inflammatory surface marker CD-206 on the CG-155-i scaffolds on days 3 and 7 post-transfection (Fig. 2L). Nuclei count assessment displayed twice as much cell nuclei on the CG-155-i scaffolds compared to the CG control on both days 3 and 7 post-transfection (Fig. 2M). Similarly, M0 macrophage cell coverage was increased 4-fold on CG-155-i scaffolds on day 7 compared to both miRNA-free and CG-Scr-i scaffolds (Fig. 2N). Analysis of CD-206 intensity, a well-established marker of M2 macrophages, exhibited a ∼1.5-fold increase in expression in cells grown in the CG-155-i scaffolds on days 3 and 7 post-transfection (Fig. 2O). Moreover, assessment of macrophage circularity on the scaffolds (Fig. 2P) –with higher circularity being associated to pro-inflammatory phenotypes– revealed a significant decrease in cells on the CG-155-i scaffolds compared to controls on both timepoints analysed, suggesting a clear phenotypic drive towards the anti-inflammatory M2 phenotype.

### 3.3. CG-155-i scaffolds elicit a regenerative response from macrophages in a pro-inflammatory environment

Having validated the ability of the CG-155-i scaffolds on the anti-inflammatory polarisation of M0 macrophages, we next assessed their ability to influence macrophages grown in a pro-inflammatory environment conditioned with lipopolysaccharide (LPS) and interferon-gamma (IFN-γ). Analysis of metabolic activity (Fig. 3A) and DNA content (Fig. 3B) of pro-inflammatory (M1) macrophages displayed a significant increase in macrophage activity and proliferation on the CG-155-i scaffolds over a 7-day period. This trend was correlated with a ∼2-fold increase in VEGF expression on day 7 post-transfection compared to control scaffolds (Fig. 3C).

**Figure 3.**
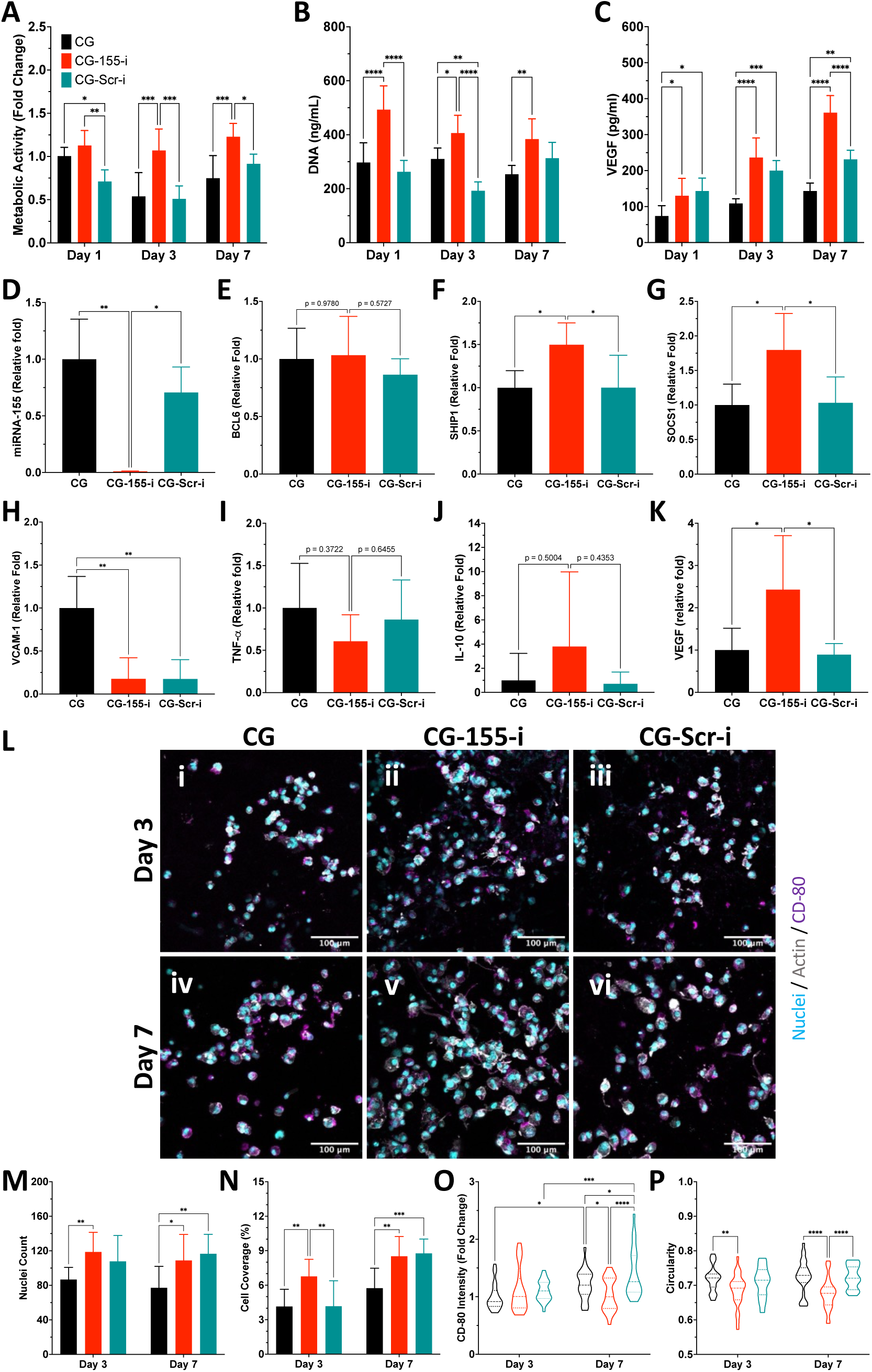
Pro-inflammatory (M1) macrophages are driven towards an anti-inflammatory (M2) phenotype on CG-155-i scaffolds. A-B) Assessment of cell viability through metabolic activity and DNA content showed increased macrophage activity and proliferation on the CG-155-i group over 7 days. C) Quantification of VEGF release was significantly enhanced with macrophages seeded on CG-155-i scaffolds. D-G) Gene expression analysis of the miRNA and directly regulated target genes SHIP1, BCL6, and SOCS1 demonstrate the activation of anti-inflammatory processes following miRNA-155 inhibition. H-K) Gene quantification of inflammatory and angiogenic-associated genes VCAM-1, TNF-α, IL-10, and VEGF showed a potential anti-inflammatory/pro-angiogenic response from M1 macrophages on CG-155-i scaffolds. L-P) Confocal visualisation of M1 macrophages on scaffolds exhibited higher macrophage count and coverage with decreased CD-80 intensity and circularity on CG-155-i scaffolds. Data shows mean ± SD (n=5), * indicates p<0.05, ** p<0.01, *** p<0.001, **** p<0.0001.

Subsequent gene expression analysis showed near elimination of the target miRNA-155 with the CG-155-i scaffolds after 3 days (Fig. 3D). This suppression did not affect the expression of the downstream target BCL6 (Fig. 3E), while expression of SHIP1 (Fig. 3F) and SOCS1 (Fig. 3G) was upregulated in accordance with what was observed with non-polarised macrophages, reaching up to ∼1.5 fold up-regulation at this timepoint compared to the miRNA-free scaffolds. Assessment of additional targets revealed a downregulation of the pro-inflammatory VCAM-1 (Fig. 3H) and TNF-α (Fig. 3I) genes, while expression of the anti-inflammatory IL-10 (Fig. 3J) and pro-angiogenic VEGF (Fig. 3K) genes were up-regulated, underscoring an anti-inflammatory response similar to what was observed with M0 macrophages despite the exposure to a more inflammatory chronic-like environment.

Following analysis of gene expression, visualisation of macrophages through confocal imaging showed that cells exhibited a reduced expression of the pro-inflammatory surface marker CD-80 (Fig. 3L) despite the increased cell number on the CG-155-i scaffolds (Fig. 3M). Quantification of cell coverage supported these results with increased coverage on days 3 and 7 post-transfection on the CG-155-i group compared to CG scaffolds (Fig. 3N). Interestingly, the cell coverage values obtained for macrophages grown on CG-Scr-i scaffolds significantly increased ∼1.5 fold compared to the CG scaffolds on day 7. Measurements of CD-80 intensity from macrophages revealed that the increased cell number and coverage was associated with a reduced expression of the surface marker on the CG-155-i scaffolds (Fig. 3O). Furthermore, these observations were supported by a reduction in circularity on both days (Fig. 3P), underscoring anti-inflammatory macrophage polarisation with lower CD-80 expression and circularity associated to the M2 macrophage phenotype.

### 3.4. Endothelial cells exposed to secretome from macrophages on CG-155-i scaffolds present a decreased expression of inflammatory proteins

Having demonstrated that scaffold-mediated miRNA-155 inhibition promotes regenerative polarisation of macrophages, the secretome profile was next analysed. Characterisation of cytokine release from M0 and M1 macrophages on miRNA-i-activated scaffolds displayed anti-inflammatory and pro-angiogenic expression profiles (Fig. 4A). The expression of pro-angiogenic growth factors, such as bFGF and VEGF, was strongly up-regulated in the CG-155-i group when compared to the miRNA-free scaffolds in the non-polarised or pro-inflammatory conditions. Importantly, these observations support previous results of VEGF release from both M0 and M1 macrophages (Fig. 2C, Fig. 3C). Moreover, expression of the ECM remodelling molecules MMP-1 and MMP-9, which are often associated with chronic inflammatory environments, were downregulated in cells grown in the CG-155-i scaffolds.

**Figure 4.**
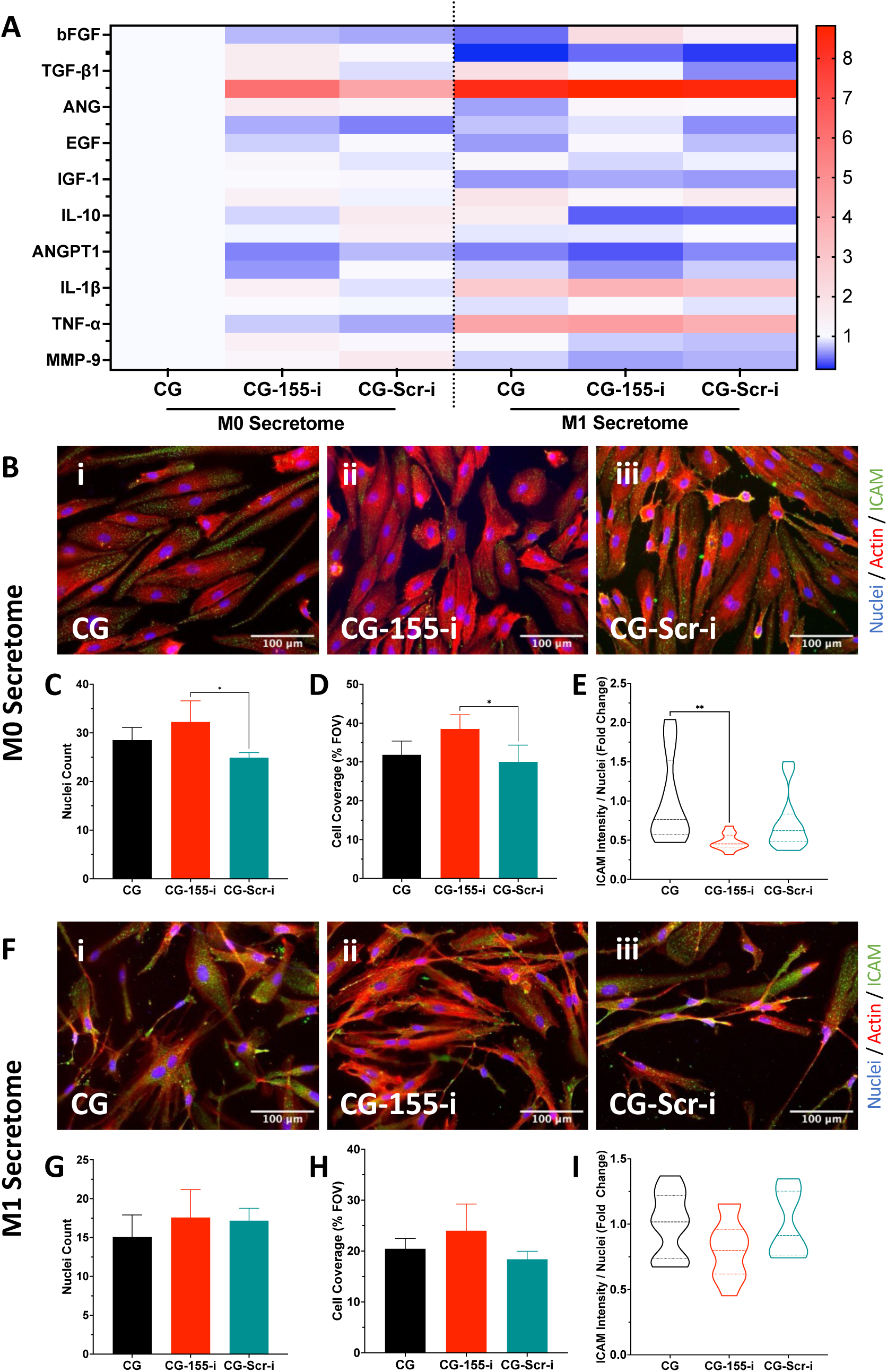
Secretome from macrophages grown on CG-155-i scaffolds induces anti-inflammatory responses on endothelial cells. A) Cytokine profile analysis revealed an increased release of pro-angiogenic and anti-inflammatory growth factors from macrophages on CG-155-i scaffolds. B-E) Endothelial cells exposed to M0 macrophage secretome show a reduced expression of pro-inflammatory ICAM in the CG-155-i group. F-I) M1 macrophage secretome on endothelial cells elicits clear morphological changes and decreased ICAM intensity in the CG-155-i group. Data shows mean ± SD (n=4), * indicates p<0.05, ** p<0.01.

In terms of inflammation-associated cytokines, the expression of interleukin-4 (IL-4, anti-inflammatory) was increased ∼5 fold in the secretome from M0 macrophages on CG-155-i scaffolds compared to the CG group. IL-4 intensity in the pro-inflammatory secretome was close to 8-fold on all groups compared to non-polarised conditions. Release of pro-inflammatory cytokines, such as IL-1α and IFN-γ and TNF-α was also reduced on miRNA-155 inhibited macrophages compared to CG and CG-Scr-i controls. However, the expression of interleukin-8 (IL-8) was upregulated with higher levels observed in the pro-inflammatory conditions.

Following the analysis of the scaffold-mediated macrophage secretome profile, modulation of inflammatory processes was investigated through macrophage secretome – endothelial cell paracrine interactions (Fig. 4, Supp. Fig. 5). Macrophage secretome-induced expression of the pro-inflammatory surface marker intercellular cell adhesion molecule (ICAM) on endothelial cells showed a clear decrease in intensity in the non-polarised conditions (Fig. 4B). Quantification of nuclei (Fig. 4C) and cell coverage (Fig. 4D) also presented an increase in the CG-155-i secretome compared to the CG-Scr-i group. Importantly, the initial visualisation of ICAM intensity was supported by a 50% decrease in ICAM expression in the CG-155-i group compared to CG group (Fig. 4E).

Similarly, when analysing ICAM expression on endothelial cells exposed to the M1 macrophage secretome, a decreasing trend was observed in the CG-155-i group compared to both CG and CG-Scr-i scaffold secretomes (Fig. 4F). Endothelial cells also presented clear morphological changes in these conditions, with the cells showing a narrower and more elongated cell body like the distinctive spindle-like shape of fibroblasts. Quantification of nuclei count (Fig. 4G) and cell coverage (Fig. 4H) also displayed a trending increased trend in the CG-155-i group compared to the miRNA-free and CG-Scr-i scaffolds. ICAM intensity measurements (Fig. 4I) exhibited a decreased expression in the CG-155-i scaffold group when compared against the CG and CG-Scr-i groups, although, these differences were not statistically significant.

### 3.5. Secretome from macrophages on CG-155-i scaffolds drive endothelial cell migration and organisation into vascular-like structures

Having validated the influence of scaffold-mediated macrophage secretome on inflammatory processes, we next assessed its effect to drive crucial angiogenic outcomes, including migration and vascular network organisation (Fig. 5, Supp. Fig. 6). Evaluation of endothelial cell migration rate through a functional assay defined apparent differences between endothelial cells exposed to M0 and M1 secretomes (Fig. 5A). In general, endothelial cells exposed to M0 secretomes demonstrated faster migration rates and complete gap closures after 24 hours compared to endothelial cells exposed to M1 secretomes, with these cells being incapable of bridging the gap at any timepoint.

**Figure 5.**
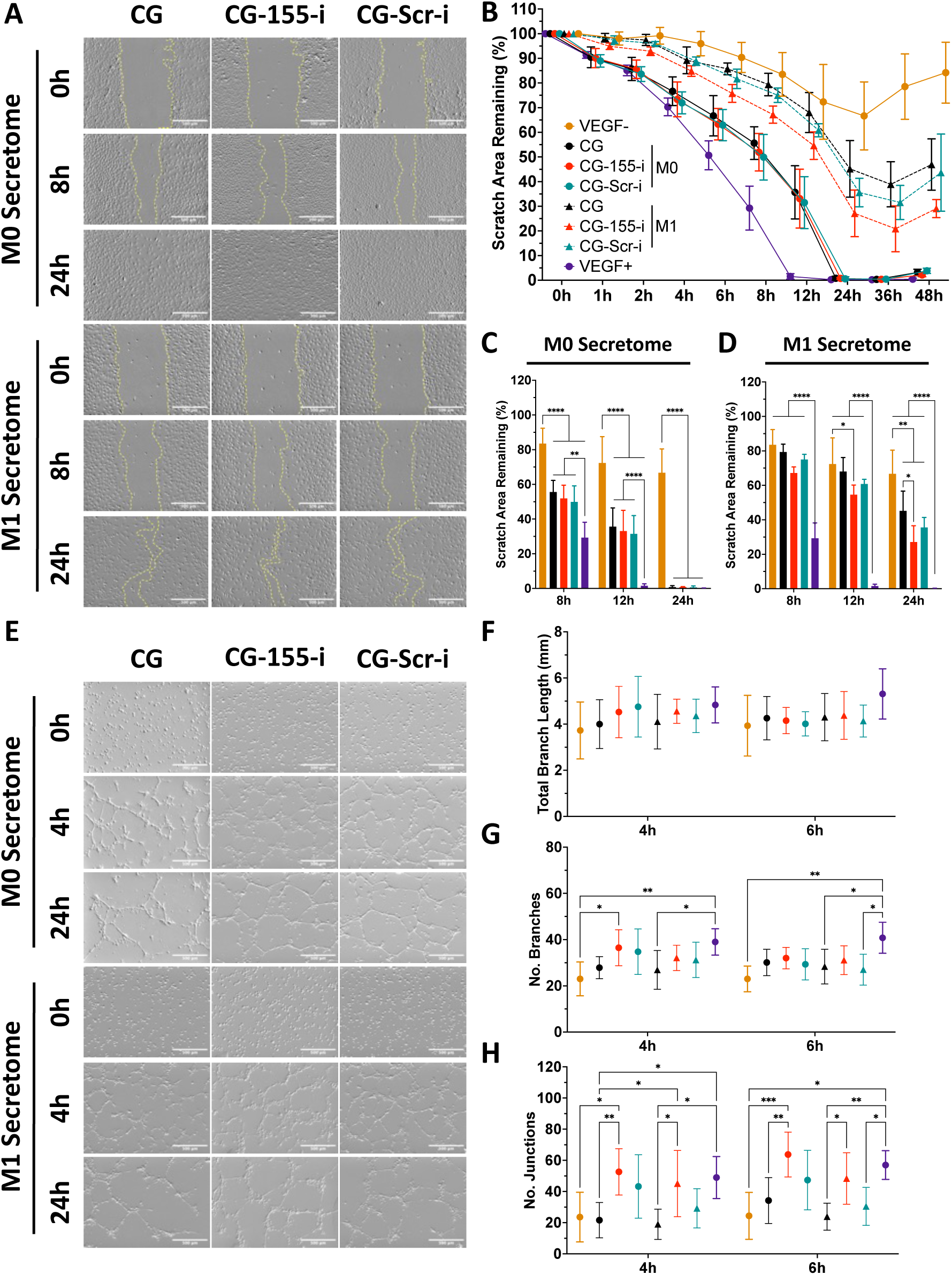
Secretome from macrophages on CG-155-i scaffolds enhances endothelial cell migration and organisation into vascular-like structures under chronic-like conditions. A) Endothelial cells exposed to M1 macrophage secretome show reduced migration rates compared to M0 conditions. B-C) Analysis of migration profiles under M0 conditions did not reveal any clear differences in behaviour between treatment groups. D-E) Endothelial cell migration rate exposed to secretome from M1 macrophages on CG-155-i scaffolds result in faster cell migration compared to the negative and miRNA-free groups after 24 hours. E) Endothelial cells show higher vascular-like organisation when exposed to M0 macrophage secretome. F-H) Secretome from CG-155-i scaffolds enables improved vascular-like complexity in both M0 and M1 conditions. Data shows mean ± SD (n=4), * indicates p<0.05, ** p<0.01, *** p>0.001, and **** p<0.0001.

Subsequent characterisation of the migration profiles confirmed our previous observations where endothelial cells exposed to CG, CG-155-i, and CG-Scr-i M0 secretomes performed within the boundaries of negative (VEGF-) and positive (VEGF+) controls (Fig. 5B). Statistical analysis of secretome-induced cell behaviour after 8, 12, and 24 hours confirmed the results that all treatment groups had greater gap closure than the negative control but failed to achieve a similar behaviour than the VEGF+ group (Fig. 5C). Importantly, assessment of migration profiles under M1 conditions showed decrease migration rates compared to M0 conditions (Fig. 5D). Endothelial cells in the CG-155-i group exhibited a faster migratory profile. This was further confirmed at 12 and 24 hours, where the CG-155-i group displayed a ∼15% increase in migration rate compared to negative and miRNA-free controls.

Following assessment of migration rate, the effect of the macrophage secretome on endothelial cell vascular-like organisation was tested using a functional tube formation assay. Imaging of the cells did not show any apparent differences after 4 to 8 hours in any of the groups. However, vascular-like structure formation was only enhanced in cells exposed to M0 secretomes compared to M1 after 24 hours (Fig. 5E). Subsequent characterisation of total branch length in the M0 and M1 conditions did not reveal any statistical difference among groups within the 4–6-hour range (Fig. 5F). Nonetheless, quantification of branch number (Fig. 5G) exhibited increased branching in the CG-155-i scaffold group when compared to the negative control after 4 hours, although this effect was stabilised after 6 hours under M0 conditions. Evaluation of junction number showed evidence of greater endothelial cell organisation in the CG-155-i group, which was found to be increased after 4 and 6 hours when compared to the negative and miRNA-free scaffold groups under M0 and M1 conditions (Fig. 5H). Taken together, these findings highlight the potential of CG-155-i-mediated macrophage polarisation to drive pro-angiogenic responses from endothelial cells through improved migration and vascular-like organisation in non-polarised and pro-inflammatory conditions.

### 3.6. CG-155-i scaffolds promote neurite outgrowth in an *ex vivo* dorsal root ganglia model of axonal regrowth

Having characterised the influence of macrophage secretome on inflammatory and angiogenic processes on endothelial cells, dorsal root ganglia (DRG) were seeded on the scaffolds to assess the effect of scaffold-mediated miRNA-155 inhibition on axonal regrowth. Imaging of DRGs culture on the miRNA-free and miRNA-i-activated scaffolds demonstrated a distinct increase in neurite number and outgrowth from DRGs grown on the CG-155-i scaffolds (Fig. 6A). Quantification of released lactate dehydrogenase (LDH) release, a well-established marker of cell stress and cytotoxicity, did not show any change across all three groups (Fig. 6B), highlighting the biocompatibility of the scaffold platform and nanoparticles to support and transfect multicellular tissues. Assessment of maximum neurite length displayed the greatest values on the CG-155-i scaffolds reaching neurites of ∼1.8 mm in length compared to ∼1.1 mm and 0.6 mm on the CG and CG-Scr-i groups, respectively (Fig. 6C). Similarly, analysis of average neurite length revealed a 2-fold increase in average neurite outgrowth on the CG-155-i scaffolds compared to other scaffold groups (Fig. 6D), underscoring the beneficial potential of scaffold-mediated miRNA-155 inhibition for to support innervation into chronic skin wounds.

**Figure 6.**
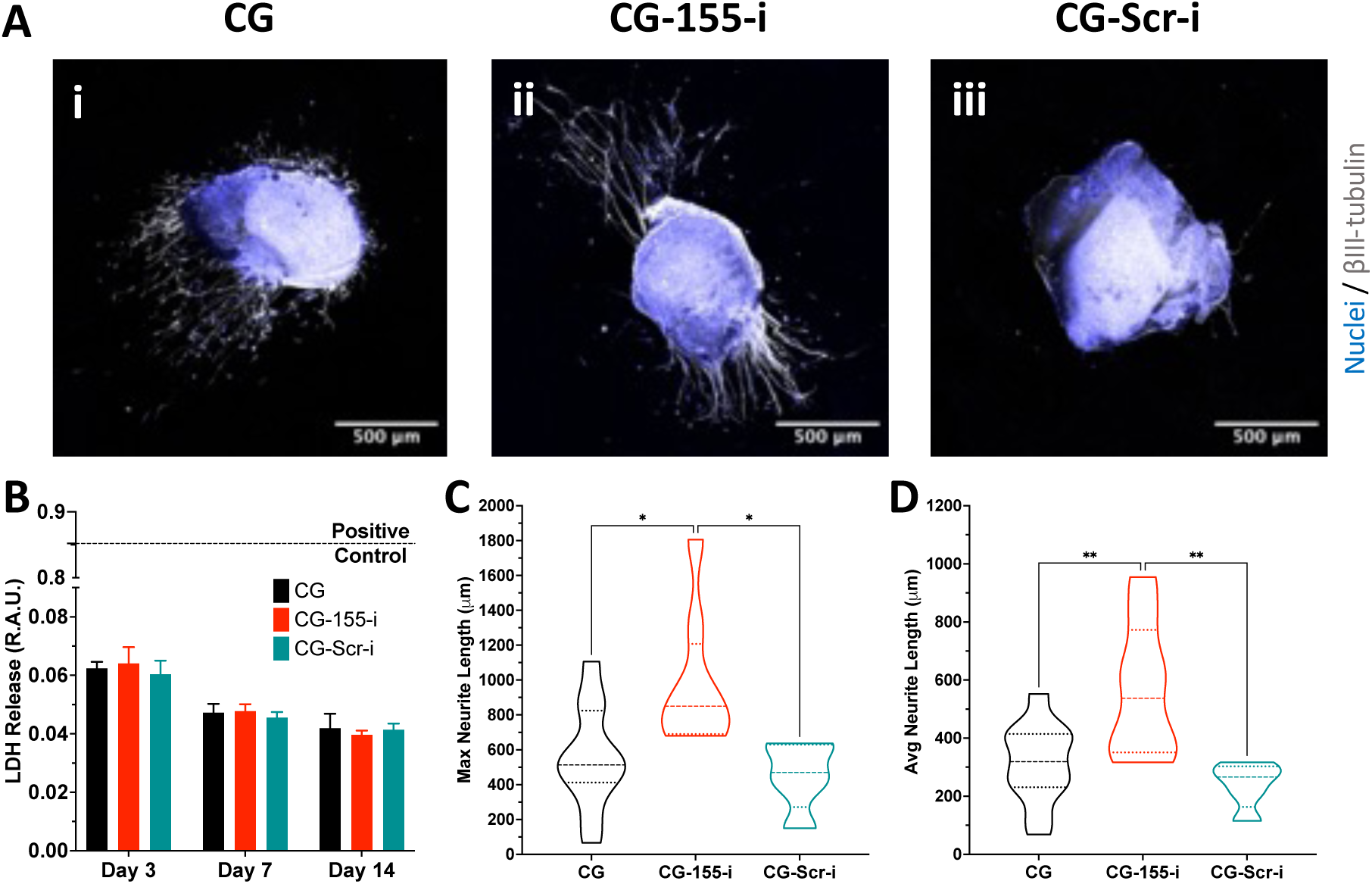
Dorsal root ganglia (DRGs) on CG-155-i scaffolds exhibit significantly improved neurite outgrowth after axonal injury. A) DRGs on CG-155-i scaffolds show higher neurite extension compared to CG and CG-Scr-i scaffolds. B) Quantification of LDH release does not display a cytotoxic response from DRGs on scaffolds. C-D) Morphological assessment of neurite length establishes the promising effects of miRNA-155 inhibition for nerve repair. Data shows mean ± SD (n=3), * indicates p<0.05, and ** p<0.01.

### 3.7. Implantation of miRNA-i-activated scaffolds demonstrates successful integration in an *in vivo* model

Following assessment of the direct and paracrine effect of scaffold-mediated miRNA-155 inhibition on inflammatory and angiogenic processes, *in vivo* compatibility post-implantation was investigated in a chicken chorioallantoic membrane (CAM) model. Visually, embryo development and membrane vascularisation were maintained among all scaffold groups with similar outcomes observed in non-treated embryos, suggesting viable integration of scaffolds (Fig. 7A). Moreover, quantification of total vessel length (Fig. 7B), average vessel length (Fig. 7C), tube number (Fig. 7D), and junction number (Fig. 7E) did not reveal any apparent differences among the different scaffold groups, indicating that scaffold implantation did not disrupt vascularisation of the membrane. Subsequent histological imaging of scaffold sections with haematoxylin and eosin (Fig. 7F) and Masson’s-Goldner trichrome (Fig. 7G) stains showed cellular infiltration, scaffold integration, and remodelling, confirming previous observations.

**Figure 7.**
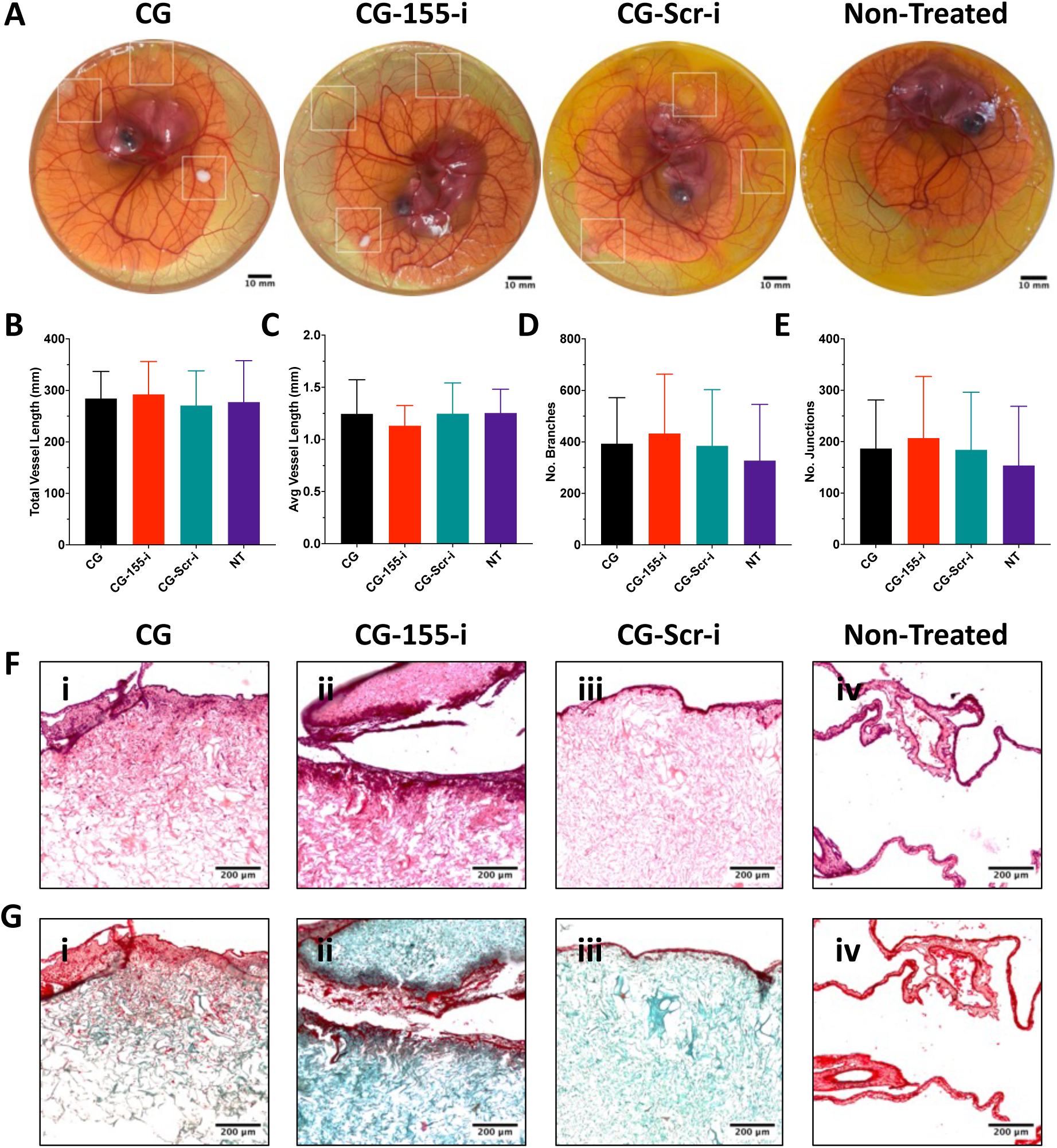
Post-implantation assessment of chicken vascularised membranes displays successful integration of miRNA-i-activated scaffolds. A) Visualisation of vasculature does not present differences in chicken embryo development. B-E) Quantification of vasculature properties do not reveal differences among scaffold-implanted embryos and non-treated embryos. F-G) Histological staining of scaffold sections showed evidence of cell infiltration and scaffold remodelling on scaffold-treated groups. Data shows mean ± SD (n=4).

## 4. Discussion

The overarching aim of this study was to develop a miRNA-155 inhibitor-activated scaffold capable of inhibiting miRNA-155 and promoting regenerative outcomes *via* reduced inflammation, enhanced vascularisation, and improved nerve regeneration for chronic wound healing applications. Initially, the physicochemical properties of miRNA-155 inhibitor complexed GET nanoparticles were characterised, followed by the formation of miRNA-155-i-activated scaffolds that supported the growth of resident dermal cells. Following this, scaffold-mediated miRNA-155 inhibition in non-polarised (M0) and pro-inflammatory (M1) macrophages displayed a phenotypic drive towards an anti-inflammatory (M2) polarisation. Confirmation of the regenerative potential of the miRNA-i-activated scaffold *via* macrophage polarisation was validated through inflammatory and angiogenic functional assays with endothelial cells. The effect of scaffold-mediated miRNA-155-i delivery to dorsal root ganglia (DRG) exhibited promising neurogenic outcomes in the form of enhanced axonal regrowth, essential for the synergistic repair of chronic wounds across the skin-nerve axis. Finally, *in vivo* assessment of miRNA-155-i-activated scaffold implantation in chicks demonstrated successful membrane integration without disruption of angiogenic processes. Taken together, this study establishes the regenerative potential of the miRNA-155-i-activated scaffolds as platforms with anti-inflammatory, pro-angiogenic, and neurogenic outcomes for chronic wound healing applications.

Initial steps to develop the platform involved characterising the properties of miRNA-155-i complexed GET nanoparticles to verify optimal size, charge, polydispersity index (PDI), complexation efficiency, and morphology that influence their release profile and distribution on CG scaffolds. Crucially, the size, PDI, and charge of the miRNA-155 nanoparticles lay within well-established parameters for successful cellular internalisation (<200 nm, >0.3, and >+10 mV, respectively), which also facilitates internalisation dynamics and endosomal escape.[56], [57], [58] Moreover, nanoparticle were mostly retained within the scaffold for up to 14 days with the bioactivity of the miRNA-155 inhibitor effectively downregulating the target miRNA for up to 7 days, suggesting the sustained therapeutic activity of the cargo from the biomaterial. Similarly, nanoparticle morphology and distribution on the CG scaffolds align with previous observations in the literature, where GET-based systems and similar peptides have been used.[24], [47], [59]

MiRNA-155 is widely recognised as a master regulator of inflammatory processes[35], with its overexpression closely associated with pathological conditions, including chronic wounds. Given its central role in driving inflammation, immune cells – particularly macrophages – are considered primary targets for miRNA-155 inhibition strategies. However, localised scaffold-mediated miRNA-155 inhibition within chronic wound environments presents a challenge in achieving cell-specific uptake, meaning other resident dermal cells may also be exposed to the therapeutic miRNA inhibitor. Crucially, *in vitro* culture of dermal fibroblasts and endothelial cells on miRNA-155-activated scaffolds did not elicit cytotoxic or adverse responses, underscoring the compatibility of the miRNA-i-activated platform across multiple skin cell types. In contrast, endothelial cells cultured on CG-Scr-i-activated scaffolds displayed reduced VE-cadherin expression. Notably, this effect was absent in the CG-155-i group, suggesting that the response was due to the scrambled miRNA sequence itself rather than an off-target action of the GET peptide.[60], [61], [62] Together, these findings underscore the biosafety of the GET peptide-CG scaffold system and support its potential as a versatile platform for the delivery of nucleic acids.

Macrophages play a central role in the pathogenesis and persistence of chronic wounds, where their dysregulated activity sustains an inflammatory microenvironment.[63] Both infiltrating monocyte-derived (M0) and tissue-resident (M1) macrophages are exposed to chronic inflammatory cues that reinforce phenotypic imbalances, particularly favouring the pro-inflammatory M1 phenotype.[64] In this context, these miRNA-i-activated scaffolds demonstrated a promising capacity to modulate macrophage behaviour. Upon exposure, both non-polarised (M0) and pro-inflammatory (M1) macrophages exhibited a shift towards an anti-inflammatory state, as evidenced by anti-inflammatory gene expression, pro-inflammatory marker CD-80 downregulation, and increased anti-inflammatory CD-206 surface marker profiles.

Mechanistically, miRNA-155 inhibition *via* the CG scaffolds led to the upregulation of critical inflammatory mediators – SOCS1 and SHIP1 – which are commonly suppressed in chronic wound environments.[35] This restoration is associated with the re-engagement of endogenous negative feedback mechanisms that suppress key inflammatory signalling pathways, including JAK/STAT[65], NF-κB[66], and PI3K/Akt[38]. As a result, anti-inflammatory signalling was enhanced through reduced TNF-α and increased IL-10 expression which is validated by previous reports in the literature.[67], [68] In parallel, the scaffolds also stimulated the release of pro-angiogenic factors such as VEGF, facilitating crosstalk with endothelial cells to promote neovascularisation. Crucially, the therapeutic relevance of this approach extends beyond cutaneous chronic wounds. Dysregulation of SOCS1 and SHIP1 has been implicated in a variety of inflammatory pathologies, including colitis[69], atherosclerosis[70], and epilepsy[71], underscoring the broader translational potential of the miRNA-155-i-activated scaffold platform in inflammation-driven diseases.

Macrophages also present a highly dynamic secretory activity, which enables them to shape the wound microenvironment through paracrine signalling.[72] Understanding the secretome profile following scaffold-mediated miRNA-155 inhibition in non-polarised M0 and pro-inflammatory M1 macrophages is essential to predict downstream cellular interactions and tissue responses. In both polarisation states, scaffold-mediated miRNA-155 inhibition upregulated key regenerative mediators, including VEGF and IL-4, both of which are associated with M2 macrophage polarisation, enhanced angiogenesis, and tissue repair. Concurrently, the expression of pro-inflammatory and matrix-degrading molecules such as IFN-γ, IL-1, and MMP-9 was significantly reduced. This shift not only supports an anti-inflammatory macrophage phenotype but also disrupts the pathological feedback loop in which pro-inflammatory cytokines drive miRNA-155 overexpression, further sustaining inflammation.[73] In particular, MMP-9 is frequently elevated in chronic wounds and, in fact, its suppression has been linked with improved regenerative outcomes.[74], [75] Moreover, the simultaneous upregulation of VEGF and IL-4 reinforces the notion that miRNA-155 inhibition effectively reprograms macrophages towards a reparative, M2 phenotype conducive to wound resolution and angiogenesis.[63], [76]

Compelling evidence of the paracrine influence exerted by macrophages was observed through the modulation of intercellular cell adhesion molecule-1 (ICAM-1) expression in endothelial cells. During the inflammatory phase of physiological wound healing, endothelial cells upregulate adhesion molecules such as P-selectin, E-selectin, VCAM-1, and ICAM-1. They mediate the adhesion and transmigration of leukocytes and monocytes across the endothelium.[77] These events are essential for immune cell recruitment and the clearance of pathogens and cellular debris. However, under pathological conditions, the failure to resolve adhesion molecule expression results in excessive immune cell infiltration and tissue damage.[77] Crucially, multiple studies have linked miRNA-155 deletion to decreased ICAM-1 expression, primarily through SOCS1-dependent inhibition of pro-inflammatory signalling.[78], [79] In line with this, endothelial cells exposed to the macrophage secretome following miRNA-155 inhibition exhibited a marked reduction in ICAM-1 expression under both non-polarised and pro-inflammatory conditions. These findings suggest that our miRNA-i-activated scaffold platform restores endothelial cell homeostasis, reduces leukocyte recruitment, and has the potential to mitigate chronic inflammation *via* macrophage-mediated paracrine signalling.

Angiogenic and vasculogenic processes are dependent on endothelial cell migration and organisation into vascular networks.[80], [81] These are chemotactically driven and require the availability of angiogenic growth factors such as VEGF and bFGF.[80] However, in chronic wound environments, these beneficial cytokines are scarce, leaving endothelial cells exposed to elevated levels of pro-inflammatory mediators that suppress angiogenic responses.[82] For instance, IFN-y upregulates ICAM-1 and VCAM-1 expression in endothelial cells and compromises endothelial barrier integrity by facilitating excessive macrophage extravasation – a hallmark of chronic wounds that inhibits proper vascularisation.[83] Crucially, scaffold-mediated inhibition of miRNA-155 in macrophages significantly enhanced the secretion of pro-angiogenic cytokines, including VEGF. This outcome then promoted endothelial cell migration and the formation of vascular-like networks, even under pro-inflammatory conditions, thereby further underscoring the multi-faceted regenerative potential of the miRNA-155-i-activated platform.

In addition to persistent inflammation and impaired angiogenesis, neuropathy represents another key but often overlooked pathological feature of chronic wounds.[84] Cutaneous peripheral nerves contribute actively to wound regeneration through bidirectional crosstalk with multiple cell types – including macrophages and endothelial cells – modulating inflammatory and angiogenic pathways.[9], [10], [11], [12] Notably, miRNA-155 inhibition in peripheral neurons has been shown to support regeneration by promoting axonal outgrowth and reducing neurotoxicity.[39], [40] Consistent with this, dorsal root ganglia (DRG) cultured on the miRNA-i-activated scaffolds exhibited enhanced βIII-tubulin-positive axon extension without any evident cytotoxicity, highlighting the scaffold’s potential to support neurogenic outcomes alongside vascular and immune repair.

The *in vivo* chicken chorioallantoic membrane (CAM) model is a widely employed preclinical platform for evaluating the biocompatibility, integration, and angiogenic potential of biomaterials.[85] While the implantation of miRNA-i-activated scaffolds into the CAM did not result in a significant increase in neovascularisation, successful integration of the miRNA-i-activated scaffolds was observed and did not disrupt the formation of healthy vasculature. This effect is likely attributable to the unique developmental timeline of the chicken embryo. While chick embryos mature rapidly – reaching full development within 21 days – their immune system develops more gradually. Innate immune cells, including macrophages, begin to appear after embryonic day 6, with functional inflammatory responses emerging around day 9.[85] Since the therapeutic efficacy of the scaffolds depend on miRNA-155 inhibition in macrophages, optimal activation of the platform would require implantation during or after this developmental stage. However, ethical considerations regarding embryonic pain perception and neural activity necessitate early termination of incubation. Studies indicate that nociception and higher brain activity become detectable around embryonic day 13.[86] To remain within ethical standards, incubation in this study was terminated by day 12, thereby limiting the scaffold’s interaction with functionally mature macrophages. Consequently, the absence of a therapeutic effect is consistent with implantation outside the optimal therapeutic window. Nonetheless, the scaffolds exhibited no adverse responses and integrated well with the membrane, supporting their suitability for *in vivo* testing in a larger mammal.

The therapeutic modulation of nucleic acids, alongside the use of CG scaffolds, has gained considerable attention as a strategy for enhancing chronic wound healing. The combination of GET peptide with CG scaffolds has previously been shown to yield superior biological outcomes compared with conventional “gold-standard” delivery vectors.[24], [47] Moreover, the biocompatibility and regenerative benefits of CG scaffolds alone have been extensively characterised.[52], [87], [88], [89] Building on this foundation, the present study investigated the potential of scaffold-mediated delivery of miRNA inhibitor/GET nanoparticles to improve regenerative outcomes in chronic wound healing via macrophage modulation, particularly given the established link between miRNA dysregulation and persistent inflammation.[90]

Notably, miRNA-146a and miRNA-223 have been widely studied in this context. MiRNA-146a is closely associated with the regulation of IL-1, TNF-α, and NF-κB signalling pathways, all of which are key drivers of the inflammatory response.[91] Further, its upregulation also suppresses pro-inflammatory cytokine production and mitigate inflammation in both *in vitro* and *in vivo* models.[92], [93] However, concerns have been raised regarding potential off-target effects, including the promotion of fibrotic responses. MiRNA-223 also plays a critical role in modulation inflammation through its effects on macrophage polarisation. Its downregulation has been linked to enhanced M1 macrophage activity, driven by increased expression of TNF receptor-associated factor 6 (TRAF6) and heightened NF-κB signalling.[94] Conversely, scaffold-based delivery of miRNA-223 has demonstrated promising results, promoting M2 polarisation and accelerating wound closure[95], and further reinforcing the therapeutic value of miRNA targeting in chronic wound environments. While these approaches offer promise, the miRNA-155-i-activated scaffold platform presents a particularly comprehensive therapeutic profile. Through targeted inhibition of miRNA-155, the platform not only modulates inflammation and promotes neurogenic processes but also supports angiogenic responses through paracrine signalling. Effectively, this multi-faceted regenerative capacity positions this miRNA-i-activated scaffold as a compelling and representative strategy for chronic wound healing applications.

## 5. Conclusion

Overall, this study outlines the development of a miRNA-155-i-activated scaffold that promotes anti-inflammatory, pro-angiogenic, and pro-neurogenic responses for potential application in chronic wound healing. The incorporation of miRNA-155 inhibitor complexed GET within collagen-GAG scaffolds enabled a wider multi-faceted therapeutic outcome with anti-inflammatory responses from non-polarised and pro-inflammatory macrophages. Angiogenic processes were enhanced through paracrine signalling with endothelial cells while dorsal root ganglia showed improved neurogenic outcomes. Moreover, *in vivo* validation of the scaffold platform displayed no cytotoxic effects and demonstrated successful integration into host tissue. Taken together, this study presents a promising alternative in the treatment of chronic wounds with the development of miRNA-155-i-activated scaffolds capable of driving regenerative outcomes.

## Author Contributions

JCPC (data curation, formal analysis, visualisation, investigation, methodology, and writing – original draft), MD (investigation, methodology, and writing – review and editing), MM (investigation, methodology, and writing – review and editing), COC (investigation, methodology, and writing – review and editing), TKM (investigation, methodology, and writing – review and editing), JM (investigation and writing – review and editing), AD (methodology, resources, and writing – review and editing), JED (resources), CJK (conceptualization, supervision, and writing – review and editing), SB (conceptualization, supervision, and writing – review and editing), FOB (conceptualization, supervision, funding acquisition, and writing – review and editing).

## Data Availability

Data for this article are openly available at Open Science Framework (OSF) Repository at https://doi.org/10.17605/OSF.IO/6HR8K under the terms of the Creative Commons Attribution 4.0 (CC-BY 4.0) license.

## Acknowledgements

The authors acknowledge the Research Ireland Advanced Materials and Bioengineering Research (AMBER) Centre for providing financial support (SFI/12/RC/2278_P2). The authors would also like to acknowledge financial support from Debra Ireland (22563A01), the Higher Education Authority (HEA), the Department of Further and Higher Education, Research, Innovation and Science (DFHERIS), the Shared Island Fund through the North South Research Program, the Health Research Board (HRB) Investigator-Led Projects (ILP) 2024 (ILP-POR-2024-064), and HORIZON EUROPE under the WIDERA Twinning Programme (Grant 101079123; ‘REGENEU’). COC, AD and FOB acknowledge funding from the Irish Rugby Football Union Charitable Trust (IRFU-CT). JED would like to acknowledge funding by the 599 Defence Accelerator (DASA, DSTL) (Award reference: ACC6007330), the European 600 Research Council under the European Community’s Seventh Framework Programme 601 (FP7/2007–2013)/ERC grant agreement 227845, the 602 Medical Research Council (grant number MR/K026682/1); the Engineering and 603 Physical Sciences Research Council; and the Biotechnology and Biological Sciences 604 Research Council, for the UK Regenerative Medicine 605 Platform Hub “Acellular Approaches for Therapeutic Delivery”. Additionally, the authors would like to acknowledge the support given by Dr Brenton Cavanagh. Collagen materials were provided by Integra Life Sciences, Inc. through a Material Transfer Agreement.

## References

[1] R. G. Frykberg and J. Banks, “Challenges in the Treatment of Chronic Wounds,” Adv. Wound Care, vol. 4, no. 9, Art. no. 9, Sept. 2015, doi: 10.1089/wound.2015.0635.

[2] P. Schilrreff and U. Alexiev, “Chronic Inflammation in Non-Healing Skin Wounds and Promising Natural Bioactive Compounds Treatment,” Int. J. Mol. Sci., vol. 23, no. 9, p. 4928, Apr. 2022, doi: 10.3390/ijms23094928.

[3] C. Lindholm and R. Searle, “Wound management for the 21st century: combining effectiveness and efficiency,” Int. Wound J., vol. 13, no. S2, pp. 5–15, July 2016, doi: 10.1111/iwj.12623.

[4] P. Gillespie, L. Carter, C. McIntosh, and G. Gethin, “Estimating the health-care costs of wound care in Ireland,” J. Wound Care, vol. 28, no. 6, pp. 324–330, June 2019, doi: 10.12968/jowc.2019.28.6.324.

[5] A. L. Laiva, F. J. O’Brien, and M. B. Keogh, “Innovations in gene and growth factor delivery systems for diabetic wound healing,” J. Tissue Eng. Regen. Med., vol. 12, no. 1, pp. e296–e312, 2018, doi: 10.1002/term.2443.

[6] R. T. Beyene, S. L. Derryberry, and A. Barbul, “The Effect of Comorbidities on Wound Healing,” Surg. Clin. North Am., vol. 100, no. 4, pp. 695–705, Aug. 2020, doi: 10.1016/j.suc.2020.05.002.

[7] H. N. Wilkinson and M. J. Hardman, “Wound healing: cellular mechanisms and pathological outcomes,” Open Biol., vol. 10, no. 9, p. 200223, Sept. 2020, doi: 10.1098/rsob.200223.

[8] F. M. Davis, A. Kimball, A. Boniakowski, and K. Gallagher, “Dysfunctional Wound Healing in Diabetic Foot Ulcers: New Crossroads,” Curr. Diab. Rep., vol. 18, no. 1, p. 2, Jan. 2018, doi: 10.1007/s11892-018-0970-z.

[9] A. P. W. Johnston and F. D. Miller, “The Contribution of Innervation to Tissue Repair and Regeneration,” Cold Spring Harb. Perspect. Biol., vol. 14, no. 9, p. a041233, Sept. 2022, doi: 10.1101/cshperspect.a041233.

[10] M. Ashrafi, M. Baguneid, and A. Bayat, “The Role of Neuromediators and Innervation in Cutaneous Wound Healing,” Acta Derm. Venereol., vol. 96, no. 5, Art. no. 5, 2016, doi: 10.2340/00015555-2321.

[11] A. G. Berger, J. J. Chou, and P. T. Hammond, “Approaches to Modulate the Chronic Wound Environment Using Localized Nucleic Acid Delivery,” Adv. Wound Care, vol. 10, no. 9, Art. no. 9, 2021, doi: 10.1089/wound.2020.1167.

[12] S. Kannan, M. Lee, S. Muthusamy, A. Blasiak, G. Sriram, and T. Cao, “Peripheral sensory neurons promote angiogenesis in neurovascular models derived from hESCs,” Stem Cell Res., vol. 52, p. 102231, Apr. 2021, doi: 10.1016/j.scr.2021.102231.

[13] L. Yan et al., “Macrophage plasticity: signaling pathways, tissue repair, and regeneration,” MedComm, vol. 5, no. 8, p. e658, Aug. 2024, doi: 10.1002/mco2.658.

[14] P. Krzyszczyk, R. Schloss, A. Palmer, and F. Berthiaume, “The Role of Macrophages in Acute and Chronic Wound Healing and Interventions to Promote Pro-wound Healing Phenotypes,” Front. Physiol., vol. 9, p. 419, May 2018, doi: 10.3389/fphys.2018.00419.

[15] R. Rios, A. Jablonka-Shariff, C. Broberg, and A. K. Snyder-Warwick, “Macrophage roles in peripheral nervous system injury and pathology: Allies in neuromuscular junction recovery,” Mol. Cell. Neurosci., vol. 111, p. 103590, Mar. 2021, doi: 10.1016/j.mcn.2021.103590.

[16] M. Sharifiaghdam, E. Shaabani, R. Faridi-Majidi, S. C. De Smedt, K. Braeckmans, and J. C. Fraire, “Macrophages as a therapeutic target to promote diabetic wound healing,” Mol. Ther., vol. 30, no. 9, pp. 2891–2908, Sept. 2022, doi: 10.1016/j.ymthe.2022.07.016.

[17] A. Hassanshahi, M. Moradzad, S. Ghalamkari, M. Fadaei, A. J. Cowin, and M. Hassanshahi, “Macrophage-Mediated Inflammation in Skin Wound Healing,” Cells, vol. 11, no. 19, p. 2953, Sept. 2022, doi: 10.3390/cells11192953.

[18] M. Li, Q. Hou, L. Zhong, Y. Zhao, and X. Fu, “Macrophage Related Chronic Inflammation in Non-Healing Wounds,” Front. Immunol., vol. 12, p. 681710, June 2021, doi: 10.3389/fimmu.2021.681710.

[19] E. Eriksson et al., “Chronic wounds: Treatment consensus,” Wound Repair Regen., vol. 30, no. 2, Art. no. 2, 2022, doi: 10.1111/wrr.12994.

[20] N. M. Vecin and R. S. Kirsner, “Skin substitutes as treatment for chronic wounds: current and future directions,” Front. Med., vol. 10, p. 1154567, Aug. 2023, doi: 10.3389/fmed.2023.1154567.

[21] Y.-F. Liu, P.-W. Ni, Y. Huang, and T. Xie, “Therapeutic strategies for chronic wound infection,” Chin. J. Traumatol., vol. 25, no. 1, pp. 11–16, Jan. 2022, doi: 10.1016/j.cjtee.2021.07.004.

[22] C. K. Sen, “Human Wound and Its Burden: Updated 2020 Compendium of Estimates,” Adv. Wound Care, vol. 10, no. 5, pp. 281–292, May 2021, doi: 10.1089/wound.2021.0026.

[23] V. R. Driver et al., “A clinical trial of Integra Template for diabetic foot ulcer treatment,” Wound Repair Regen., vol. 23, no. 6, pp. 891–900, 2015, doi: 10.1111/wrr.12357.

[24] J. C. Palomeque-Chávez et al., “Development of a VEGF-activated scaffold with enhanced angiogenic and neurogenic properties for chronic wound healing applications,” Biomater. Sci., vol. 13, pp. 1993–2011, 2025, doi: 10.1039/D4BM01051E.

[25] A. L. Laiva, F. J. O’Brien, and M. B. Keogh, “Dual delivery gene-activated scaffold directs fibroblast activity and keratinocyte epithelization,” APL Bioeng., vol. 8, no. 1, Art. no. 1, Mar. 2024, doi: 10.1063/5.0174122.

[26] T. Weng et al., “Dual gene-activated dermal scaffolds regulate angiogenesis and wound healing by mediating the coexpression of VEGF and angiopoietin-1,” Bioeng. Transl. Med., vol. 8, no. 5, Art. no. 5, Sept. 2023, doi: 10.1002/btm2.10562.

[27] D. Lou, Y. Luo, Q. Pang, W. Q. Tan, and L. Ma, “Gene-activated dermal equivalents to accelerate healing of diabetic chronic wounds by regulating inflammation and promoting angiogenesis,” Bioact. Mater., vol. 5, no. 3, Art. no. 3, 2020, doi: 10.1016/j.bioactmat.2020.04.018.

[28] J. Wang et al., “A dual gene-activated dermal scaffolds loaded with nanocomposite particles expressing of VEGF and aFGF: Promoting wound healing by early vascularization,” Int. J. Biol. Macromol., vol. 307, p. 141831, May 2025, doi: 10.1016/j.ijbiomac.2025.141831.

[29] J. C. Palomeque Chávez, M. McGrath, C. J. Kearney, S. Browne, and F. J. O’Brien, “Biomaterial scaffold-based gene delivery for the repair of complex wounds: Challenges, progress, and future perspectives,” Cell Biomater., vol. 1, no. 6, p. 100073, Apr. 2025, doi: 10.1016/j.celbio.2025.100073.

[30] E. Bayraktar, R. Bayraktar, H. Oztatlici, G. Lopez-Berestein, P. Amero, and C. Rodriguez-Aguayo, “Targeting miRNAs and Other Non-Coding RNAs as a Therapeutic Approach: An Update,” Non-Coding RNA, vol. 9, no. 2, p. 27, Apr. 2023, doi: 10.3390/ncrna9020027.

[31] D. Aljamal, P. S. Iyengar, and T. T. Nguyen, “Translational Challenges in Drug Therapy and Delivery Systems for Treating Chronic Lower Extremity Wounds,” Pharmaceutics, vol. 16, no. 6, p. 750, June 2024, doi: 10.3390/pharmaceutics16060750.

[32] M. G. Monaghan, R. Borah, C. Thomsen, and S. Browne, “Thou shall not heal: Overcoming the non-healing behaviour of diabetic foot ulcers by engineering the inflammatory microenvironment,” Adv. Drug Deliv. Rev., vol. 203, p. 115120, Dec. 2023, doi: 10.1016/j.addr.2023.115120.

[33] S. Pasca, A. Jurj, B. Petrushev, C. Tomuleasa, and D. Matei, “MicroRNA-155 Implication in M1 Polarization and the Impact in Inflammatory Diseases,” Front. Immunol., vol. 11, p. 625, Apr. 2020, doi: 10.3389/fimmu.2020.00625.

[34] J. Ye, Y. Kang, X. Sun, P. Ni, M. Wu, and S. Lu, “MicroRNA-155 Inhibition Promoted Wound Healing in Diabetic Rats,” Int. J. Low. Extrem. Wounds, vol. 16, no. 2, Art. no. 2, 2017, doi: 10.1177/1534734617706636.

[35] G. Mahesh and R. Biswas, “MicroRNA-155: A Master Regulator of Inflammation,” J. Interferon Cytokine Res., vol. 39, no. 6, Art. no. 6, June 2019, doi: 10.1089/jir.2018.0155.

[36] E. Koumpis et al., “The Role of microRNA-155 as a Biomarker in Diffuse Large B-Cell Lymphoma,” Biomedicines, vol. 12, no. 12, p. 2658, Nov. 2024, doi: 10.3390/biomedicines12122658.

[37] K. Kopp et al., “STAT5-mediated expression of oncogenic miR-155 in cutaneous T-cell lymphoma,” Cell Cycle, vol. 12, no. 12, pp. 1939–1947, June 2013, doi: 10.4161/cc.24987.

[38] X. Huang et al., “Quantitative Proteomics Reveals that miR-155 Regulates the PI3K-AKT Pathway in Diffuse Large B-Cell Lymphoma,” Am. J. Pathol., vol. 181, no. 1, pp. 26–33, July 2012, doi: 10.1016/j.ajpath.2012.03.013.

[39] J. Chen, C. Li, W. Liu, B. Yan, X. Hu, and F. Yang, “miRNA-155 silencing reduces sciatic nerve injury in diabetic peripheral neuropathy,” J. Mol. Endocrinol., vol. 63, no. 3, pp. 227–238, Oct. 2019, doi: 10.1530/JME-19-0067.

[40] A. D. Gaudet et al., “miR-155 Deletion in Mice Overcomes Neuron-Intrinsic and Neuron-Extrinsic Barriers to Spinal Cord Repair,” J. Neurosci., vol. 36, no. 32, pp. 8516–8532, Aug. 2016, doi: 10.1523/JNEUROSCI.0735-16.2016.

[41] K. Obora et al., “Inflammation-induced miRNA-155 inhibits self-renewal of neural stem cells via suppression of CCAAT/enhancer binding protein β (C/EBPβ) expression,” Sci. Rep., vol. 7, no. 1, p. 43604, Feb. 2017, doi: 10.1038/srep43604.

[42] E. D. Koval et al., “Method for widespread microRNA-155 inhibition prolongs survival in ALS-model mice,” Hum. Mol. Genet., vol. 22, no. 20, pp. 4127–4135, Oct. 2013, doi: 10.1093/hmg/ddt261.

[43] Ö. Tezgel et al., “Collagen scaffold-mediated delivery of NLC/siRNA as wound healing materials,” J. Drug Deliv. Sci. Technol., vol. 55, no. August 2019, 2020, doi: 10.1016/j.jddst.2019.101421.

[44] A. I. S. Van Den Berg, C.-O. Yun, R. M. Schiffelers, and W. E. Hennink, “Polymeric delivery systems for nucleic acid therapeutics: Approaching the clinic,” J. Controlled Release, vol. 331, pp. 121–141, Mar. 2021, doi: 10.1016/j.jconrel.2021.01.014.

[45] R. M. Raftery, E. G. Tierney, C. M. Curtin, S. A. Cryan, and F. J. O’Brien, “Development of a gene-activated scaffold platform for tissue engineering applications using chitosan-pDNA nanoparticles on collagen-based scaffolds,” J. Controlled Release, vol. 210, pp. 84–94, 2015, doi: 10.1016/j.jconrel.2015.05.005.

[46] H. A. D. M. Abu-Awwad, L. Thiagarajan, and J. E. Dixon, “Controlled release of GAG-binding enhanced transduction (GET) peptides for sustained and highly efficient intracellular delivery,” Acta Biomater., vol. 57, pp. 225–237, 2017, doi: 10.1016/j.actbio.2017.04.028.

[47] R. M. Raftery et al., “Highly versatile cell-penetrating peptide loaded scaffold for efficient and localised gene delivery to multiple cell types: From development to application in tissue engineering,” Biomaterials, vol. 216, no. June, 2019, doi: 10.1016/j.biomaterials.2019.119277.

[48] R. N. Power, B. L. Cavanagh, J. E. Dixon, C. M. Curtin, and F. J. O’Brien, “Development of a Gene-Activated Scaffold Incorporating Multifunctional Cell-Penetrating Peptides for pSDF-1α Delivery for Enhanced Angiogenesis in Tissue Engineering Applications,” Int. J. Mol. Sci., vol. 23, no. 3, Art. no. 3, Jan. 2022, doi: 10.3390/ijms23031460.

[49] J. E. Dixon et al., “Highly efficient delivery of functional cargoes by the synergistic effect of GAG binding motifs and cell-penetrating peptides,” Proc. Natl. Acad. Sci. U. S. A., vol. 113, no. 3, Art. no. 3, 2016, doi: 10.1073/pnas.1518634113.

[50] H. M. Eltaher, J. Yang, K. M. Shakesheff, and J. E. Dixon, “Highly efficient intracellular transduction in three-dimensional gradients for programming cell fate,” Acta Biomater., vol. 41, pp. 181–192, Sept. 2016, doi: 10.1016/j.actbio.2016.06.004.

[51] J. Schindelin et al., “Fiji: an open-source platform for biological-image analysis,” Nat. Methods, vol. 9, no. 7, pp. 676–682, July 2012, doi: 10.1038/nmeth.2019.

[52] M. McGrath et al., “A Biomimetic, Bilayered Antimicrobial Collagen-Based Scaffold for Enhanced Healing of Complex Wound Conditions,” ACS Appl. Mater. Interfaces, vol. 15, no. 14, pp. 17444–17458, Apr. 2023, doi: 10.1021/acsami.2c18837.

[53] J. Maughan et al., “Collagen/pristine graphene as an electroconductive interface material for neuronal medical device applications,” *Appl*. Mater. Today, vol. 29, p. 101629, Dec. 2022, doi: 10.1016/j.apmt.2022.101629.

[54] I. Woods et al., “Biomimetic Scaffolds for Spinal Cord Applications Exhibit Stiffness-Dependent Immunomodulatory and Neurotrophic Characteristics,” Adv. Healthc. Mater., vol. 11, no. 3, pp. 1–17, 2022, doi: 10.1002/adhm.202101663.

[55] G. Carpentier et al., “Angiogenesis Analyzer for ImageJ — A comparative morphometric analysis of ‘Endothelial Tube Formation Assay’ and ‘Fibrin Bead Assay,’” *Sci*. Rep., vol. 10, no. 1, p. 11568, July 2020, doi: 10.1038/s41598-020-67289-8.

[56] S. Mazumdar, D. Chitkara, and A. Mittal, “Exploration and insights into the cellular internalization and intracellular fate of amphiphilic polymeric nanocarriers,” Acta Pharm. Sin. B, vol. 11, no. 4, pp. 903–924, Apr. 2021, doi: 10.1016/j.apsb.2021.02.019.

[57] J. Rejman, V. Oberle, I. S. Zuhorn, and D. Hoekstra, “Size-dependent internalization of particles via the pathways of clathrin-and caveolae-mediated endocytosis,” Biochem. J., vol. 377, no. 1, pp. 159–169, Jan. 2004, doi: 10.1042/bj20031253.

[58] M. Danaei et al., “Impact of Particle Size and Polydispersity Index on the Clinical Applications of Lipidic Nanocarrier Systems,” Pharmaceutics, vol. 10, no. 2, p. 57, May 2018, doi: 10.3390/pharmaceutics10020057.

[59] J. M. Sadowska et al., “Development of miR-26a-activated scaffold to promote healing of critical-sized bone defects through angiogenic and osteogenic mechanisms,” Biomaterials, vol. 303, p. 122398, Dec. 2023, doi: 10.1016/j.biomaterials.2023.122398.

[60] H. Seok, H. Lee, E.-S. Jang, and S. W. Chi, “Evaluation and control of miRNA-like off-target repression for RNA interference,” Cell. Mol. Life Sci., vol. 75, no. 5, pp. 797–814, Mar. 2018, doi: 10.1007/s00018-017-2656-0.

[61] R. Bartoszewski and A. F. Sikorski, “Editorial focus: understanding off-target effects as the key to successful RNAi therapy,” Cell. Mol. Biol. Lett., vol. 24, no. 1, p. 69, Dec. 2019, doi: 10.1186/s11658-019-0196-3.

[62] S. Saxena, Z. O. Jónsson, and A. Dutta, “Small RNAs with Imperfect Match to Endogenous mRNA Repress Translation,” J. Biol. Chem., vol. 278, no. 45, pp. 44312–44319, Nov. 2003, doi: 10.1074/jbc.M307089200.

[63] S. Chen et al., “Macrophages in immunoregulation and therapeutics,” Signal Transduct. Target. Ther., vol. 8, no. 1, p. 207, May 2023, doi: 10.1038/s41392-023-01452-1.

[64] K. E. Martin and A. J. García, “Macrophage phenotypes in tissue repair and the foreign body response: Implications for biomaterial-based regenerative medicine strategies,” Acta Biomater., vol. 133, pp. 4–16, Oct. 2021, doi: 10.1016/j.actbio.2021.03.038.

[65] J. J. O’Shea and P. J. Murray, “Cytokine Signaling Modules in Inflammatory Responses,” Immunity, vol. 28, no. 4, pp. 477–487, Apr. 2008, doi: 10.1016/j.immuni.2008.03.002.

[66] G. Markopoulos et al., “Roles of NF-κB Signaling in the Regulation of miRNAs Impacting on Inflammation in Cancer,” Biomedicines, vol. 6, no. 2, p. 40, Mar. 2018, doi: 10.3390/biomedicines6020040.

[67] I. Prieto et al., “A mutual regulatory loop between miR-155 and SOCS1 influences renal inflammation and diabetic kidney disease,” Mol. Ther. Nucleic Acids, vol. 34, p. 102041, Dec. 2023, doi: 10.1016/j.omtn.2023.102041.

[68] D. Yee, K. M. Shah, M. C. Coles, T. V. Sharp, and D. Lagos, “MicroRNA-155 induction via TNF-α and IFN-γ suppresses expression of programmed death ligand-1 (PD-L1) in human primary cells,” J. Biol. Chem., vol. 292, no. 50, pp. 20683–20693, Dec. 2017, doi: 10.1074/jbc.M117.809053.

[69] S. Pathak et al., “MiR-155 modulates the inflammatory phenotype of intestinal myofibroblasts by targeting SOCS1 in ulcerative colitis,” Exp. Mol. Med., vol. 47, no. 5, pp. e164–e164, May 2015, doi: 10.1038/emm.2015.21.

[70] J. Ye, R. Guo, Y. Shi, F. Qi, C. Guo, and L. Yang, “miR-155 Regulated Inflammation Response by the SOCS1-STAT3-PDCD4 Axis in Atherogenesis,” Mediators Inflamm., vol. 2016, pp. 1–14, 2016, doi: 10.1155/2016/8060182.

[71] W. Yang et al., “CRYAA activates the SIRT1-PI3K/AKT signaling pathway by suppressing miR-155-5p to protect the RPE,” Arch. Biochem. Biophys., p. 110435, May 2025, doi: 10.1016/j.abb.2025.110435.

[72] S. Alivernini et al., “MicroRNA-155—at the Critical Interface of Innate and Adaptive Immunity in Arthritis,” Front. Immunol., vol. 8, p. 1932, Jan. 2018, doi: 10.3389/fimmu.2017.01932.

[73] S. Khayati, S. Dehnavi, M. Sadeghi, J. Tavakol Afshari, S.-A. Esmaeili, and M. Mohammadi, “The potential role of miRNA in regulating macrophage polarization,” Heliyon, vol. 9, no. 11, p. e21615, Nov. 2023, doi: 10.1016/j.heliyon.2023.e21615.

[74] M. Hariono, S. H. Yuliani, E. P. Istyastono, F. D. O. Riswanto, and C. F. Adhipandito, “Matrix metalloproteinase 9 (MMP9) in wound healing of diabetic foot ulcer: Molecular target and structure-based drug design,” Wound Med., vol. 22, pp. 1–13, Sept. 2018, doi: 10.1016/j.wndm.2018.05.003.

[75] L. P. Yan et al., “Collagen/GAG scaffolds activated by RALA-siMMP-9 complexes with potential for improved diabetic foot ulcer healing,” Mater. Sci. Eng. C, 2020, doi: 10.1016/j.msec.2020.111022.

[76] C. Stockmann et al., “A Wound Size–Dependent Effect of Myeloid Cell–Derived Vascular Endothelial Growth Factor on Wound Healing,” J. Invest. Dermatol., vol. 131, no. 3, pp. 797–801, Mar. 2011, doi: 10.1038/jid.2010.345.

[77] J. Kalucka, L. Bierhansl, B. Wielockx, P. Carmeliet, and G. Eelen, “Interaction of endothelial cells with macrophages—linking molecular and metabolic signaling,” Pflüg. Arch. - Eur. J. Physiol., vol. 469, no. 3–4, pp. 473–483, Apr. 2017, doi: 10.1007/s00424-017-1946-6.

[78] H. Zhang et al., “Genistein Protects Against Ox-LDL-Induced Inflammation Through MicroRNA-155/SOCS1-Mediated Repression of NF-ĸB Signaling Pathway in HUVECs,” Inflammation, vol. 40, no. 4, pp. 1450–1459, Aug. 2017, doi: 10.1007/s10753-017-0588-3.

[79] J. Keck et al., “The Role of MicroRNA-155 in Chlamydia muridarum Infected lungs,” Microbes Infect., vol. 22, no. 8, pp. 360–365, Sept. 2020, doi: 10.1016/j.micinf.2020.02.005.

[80] L. Lamalice, F. Le Boeuf, and J. Huot, “Endothelial Cell Migration During Angiogenesis,” Circ. Res., vol. 100, no. 6, Art. no. 6, Mar. 2007, doi: 10.1161/01.RES.0000259593.07661.1e.

[81] A. Czirok, “Endothelial cell motility, coordination and pattern formation during vasculogenesis,” *WIREs Syst*. Biol. Med., vol. 5, no. 5, pp. 587–602, Sept. 2013, doi: 10.1002/wsbm.1233.

[82] T. Velnar and L. Gradisnik, “Tissue Augmentation in Wound Healing: the Role of Endothelial and Epithelial Cells,” Med. Arch., vol. 72, no. 6, p. 444, 2018, doi: 10.5455/medarh.2018.72.444-448.

[83] N. Wettschureck, B. Strilic, and S. Offermanns, “Passing the Vascular Barrier: Endothelial Signaling Processes Controlling Extravasation,” Physiol. Rev., vol. 99, no. 3, pp. 1467–1525, July 2019, doi: 10.1152/physrev.00037.2018.

[84] P. W. Ackermann and D. A. Hart, “Influence of Comorbidities: Neuropathy, Vasculopathy, and Diabetes on Healing Response Quality,” Adv. Wound Care, vol. 2, no. 8, pp. 410–421, Oct. 2013, doi: 10.1089/wound.2012.0437.

[85] M. Dhayer, A. Jordao, S. Dekiouk, D. Cleret, N. Germain, and P. Marchetti, “Implementing Chicken Chorioallantoic Membrane Assays for Validating Biomaterials in Tissue Engineering: Rationale and Methods,” J. Biomed. Mater. Res. B Appl. Biomater., vol. 112, no. 11, p. e35496, Nov. 2024, doi: 10.1002/jbm.b.35496.

[86] S. Kollmansperger et al., “Nociception in Chicken Embryos, Part II: Embryonal Development of Electroencephalic Neuronal Activity In Ovo as a Prerequisite for Nociception,” Animals, vol. 13, no. 18, p. 2839, Sept. 2023, doi: 10.3390/ani13182839.

[87] C. M. Murphy, M. G. Haugh, and F. J. O’Brien, “The effect of mean pore size on cell attachment, proliferation and migration in collagen–glycosaminoglycan scaffolds for bone tissue engineering,” Biomaterials, vol. 31, no. 3, pp. 461–466, Jan. 2010, doi: 10.1016/j.biomaterials.2009.09.063.

[88] F. J. O’Brien, B. A. Harley, I. V. Yannas, and L. J. Gibson, “The effect of pore size on cell adhesion in collagen-GAG scaffolds,” Biomaterials, vol. 26, no. 4, Art. no. 4, Feb. 2005, doi: 10.1016/j.biomaterials.2004.02.052.

[89] M. G. Haugh, C. M. Murphy, R. C. McKiernan, C. Altenbuchner, and F. J. O’Brien, “Crosslinking and Mechanical Properties Significantly Influence Cell Attachment, Proliferation, and Migration Within Collagen Glycosaminoglycan Scaffolds,” Tissue Eng. Part A, vol. 17, no. 9–10, pp. 1201–1208, May 2011, doi: 10.1089/ten.tea.2010.0590.

[90] B. Chatterjee et al., “MicroRNAs: Key modulators of inflammation-associated diseases,” Semin. Cell Dev. Biol., vol. 154, pp. 364–373, Feb. 2024, doi: 10.1016/j.semcdb.2023.01.009.

[91] X. Zhou et al., “The bone mesenchymal stem cell-derived exosomal miR-146a-5p promotes diabetic wound healing in mice via macrophage M1/M2 polarization,” Mol. Cell. Endocrinol., vol. 579, p. 112089, Jan. 2024, doi: 10.1016/j.mce.2023.112089.

[92] L. C. Dewberry et al., “Cerium oxide nanoparticle conjugation to microRNA-146a mechanism of correction for impaired diabetic wound healing,” *Nanomedicine Nanotechnol*. Biol. Med., vol. 40, p. 102483, 2022, doi: 10.1016/j.nano.2021.102483.

[93] Q. Li et al., “MiR146a-loaded engineered exosomes released from silk fibroin patch promote diabetic wound healing by targeting IRAK1,” Signal Transduct. Target. Ther., vol. 8, no. 1, p. 62, Feb. 2023, doi: 10.1038/s41392-022-01263-w.

[94] G. Zhuang et al., “A Novel Regulator of Macrophage Activation: miR-223 in Obesity-Associated Adipose Tissue Inflammation,” Circulation, vol. 125, no. 23, pp. 2892–2903, June 2012, doi: 10.1161/circulationaha.111.087817.

[95] B. Saleh et al., “Local Immunomodulation Using an Adhesive Hydrogel Loaded with miRNA-Laden Nanoparticles Promotes Wound Healing,” Small, vol. 15, no. 36, pp. 1–15, 2019, doi: 10.1002/smll.201902232.

