## Supplementary Information for "Scaffold-mediated miRNA-155 inhibition promotes regenerative macrophage polarisation leading to anti-inflammatory, angiogenic and neurogenic responses for wound healing"

### Supplementary Methods

**Supplementary Method 1.** Haematoxylin and Eosin (H&E) staining of scaffold sections.

A. Deparaffinisation

| Step | Chemical | Time |
|---|---|---|
| **1** | Xylene | 10 min |
| **2** | 100% Ethanol | 5 min |
| **3** | 95% Ethanol | 5 min |
| **4** | 70% Ethanol | 5 min |
| **5** | 50% Ethanol | 5 min |
| **6** | Tap $H_2O$ | 5 min |

B. Staining

| Step | Chemical | Time |
|---|---|---|
| **1** | Harris Haematoxylin | 5 min |

| | | |
|---|---|---|
| **2** | Running Tap $H_2O$ | 10 min |
| **3** | Acid Alcohol (0.25%HCl in 70% Ethanol) | 3 dips |
| **4** | Tap $H_2O$ | 5 dips |
| **5** | Eosin | 3 min |

C. Dehydration

| Step | Chemical | Time |
|---|---|---|
| **1** | 95% Ethanol | 5 dips |
| **2** | 100% Ethanol | 5 dips |
| **3** | Xylene | 6 min |

**Supplementary Method 2.** Masson-Goldner staining of scaffold sections.

A. Deparaffinisation

| Step | Chemical | Time |
|---|---|---|
| **1** | Xylene | 10 min |
| **2** | 100% Ethanol | 5 min |
| **3** | 95% Ethanol | 5 min |
| **4** | 70% Ethanol | 5 min |
| **5** | 50% Ethanol | 5 min |
| **6** | Tap $H_2O$ | 5 min |

### B. Staining

| Step | Chemical | Time |
|---|---|---|
| **1** | Weigert's haematoxylin | 5 min |
| **2** | Running $H_2O$ | 5 min |
| **3** | Acetic Acid 1% | 30 s |
| **4** | Reagent 1 (Azophloxine) | 10 min |
| **5** | Acetic Acid 1% | 30 s |
| **6** | Reagent 2 (Tungstophosphoric acid orange G) | 1 min |
| **7** | Acetic Acid 1% | 30 s |
| **8** | Reagent 3 (Light Green SF) | 2 min |
| **9** | Acetic Acid 1% | 30 s |

### C. Dehydration

| Step | Chemical | Time |
|---|---|---|
| **1** | 70% Ethanol | 30 s |
| **2** | 95% Ethanol | 30 s |
| **3** | 100% Ethanol | 30 s |
| **4** | Xylene | 5 min |

### Supplementary Tables

**Supplementary Table 1.** Lists of genes analysed by qRT-PCR.

| Target Protein | Target Gene Reference | GeneGlobeID |
|---|---|---|
| **18S Ribosomal RNA (18S)** | Hs_RRN18S_1_SG | QT00199367 |
| **Vascular Endothelial Growth Factor (VEGF)** | Hs_VEGFA_1_SG | QT01010184 |
| **Interleukin-10 (IL-10)** | Hs_IL10_1_SG | QT00041685 |
| **Tumour Necrosis Factor-α** | Hs_TNF_1_SG | QT00029162 |
| **Vascular Cell Adhesion Molecule (VCAM1)** | Hs_VCAM1_1_SG | QT00018347 |
| **Src homology 2 domains containing inositol polyphosphatase 5-phosphatase 1 (SHIP1)** | Hs_INPP5D_1_SG | QT00048370 |
| **B-cell lymphoma 6 (BCL6)** | Hs_BCL6_1_SG | QT00079233 |
| **Suppresor of cytokine signaling 1 (SOCS1)** | Hs_SOCS1_1_SG | QT00202475 |
| **microRNA-155** | hsa-miR-155-5p | 483064_mir |

### Supplementary Figures

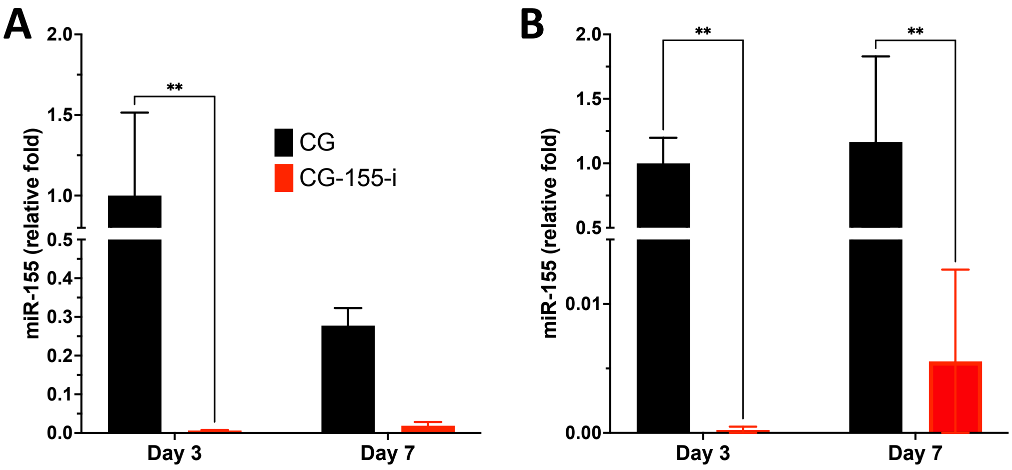


**Supplementary Figure 1. MiRNA-155 inhibition is sustained for up to 7 days following scaffold-mediated miRNA-155-i delivery in both M0 and M1 conditions**. A) Scaffold-mediated miRNA-155 inhibition in non-polarised (M0) macrophages. B) Scaffold-mediated miRNA-155 inhibition in pro-inflammatory (M1) macrophages. Data shows mean ± SD, ** indicates $p<0.01$.

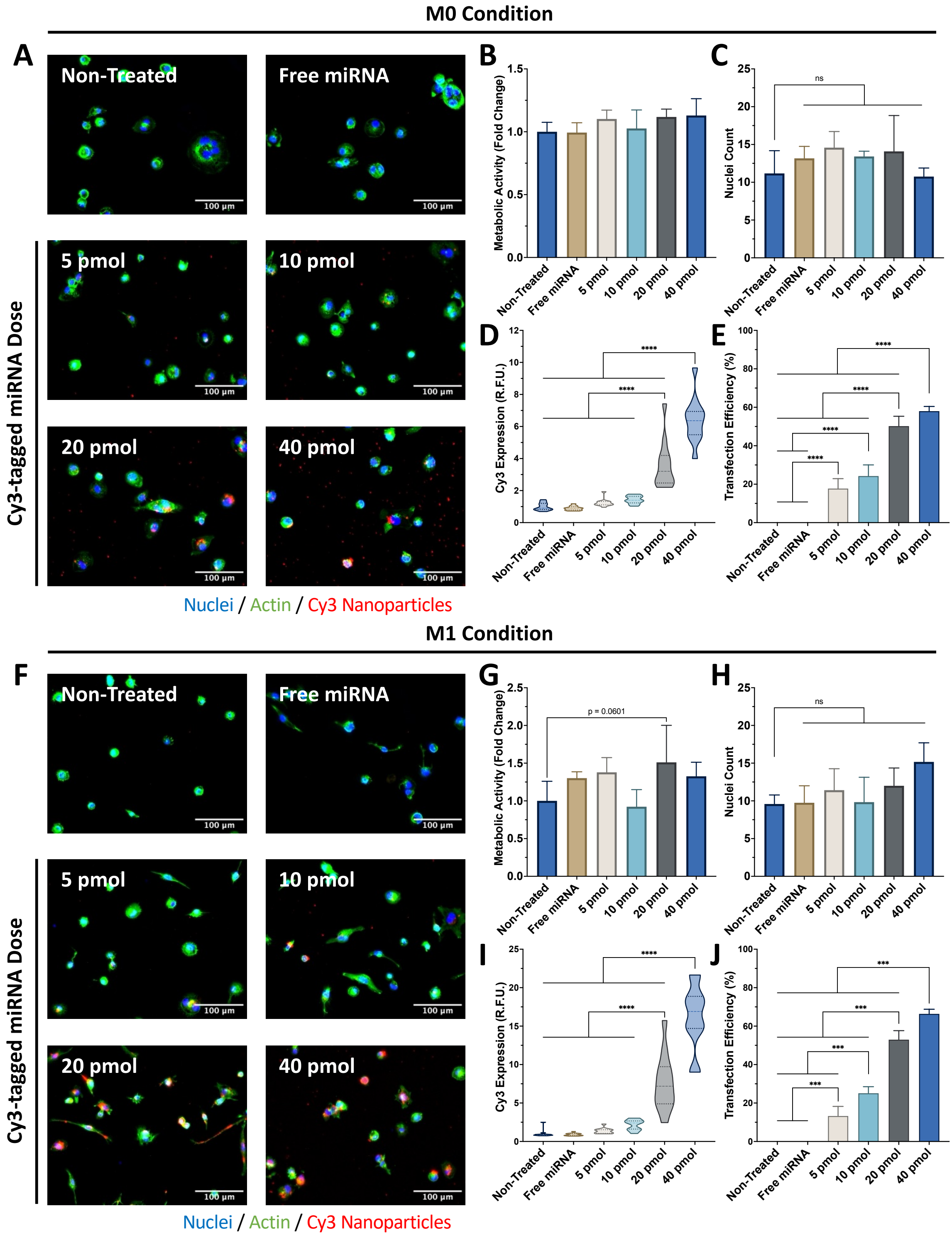


**Supplementary Figure 2. Increasing doses of miRNA-155-i nanoparticles enhance transfection efficiency in both M0 and M1 conditions**. A-C) Delivery of up to 40 pmol of Cy3-tagged miRNA-i nanoparticles does not significantly affect cell viability or number of M0 macrophages. D-E) Increasing nanoparticle dosage leads to enhanced Cy3 expression and transfection efficiency in M0 macrophages. F-H) Delivery of up to 40 pmol of Cy3-tagged miRNA-i nanoparticles does not significantly affect cell viability or number of M1 macrophages.

D-E) Increasing nanoparticle dosage leads to enhanced Cy3 expression and transfection efficiency in M1 macrophages. Data shows mean ± SD, *** indicates p<0.001, **** p<0.0001.

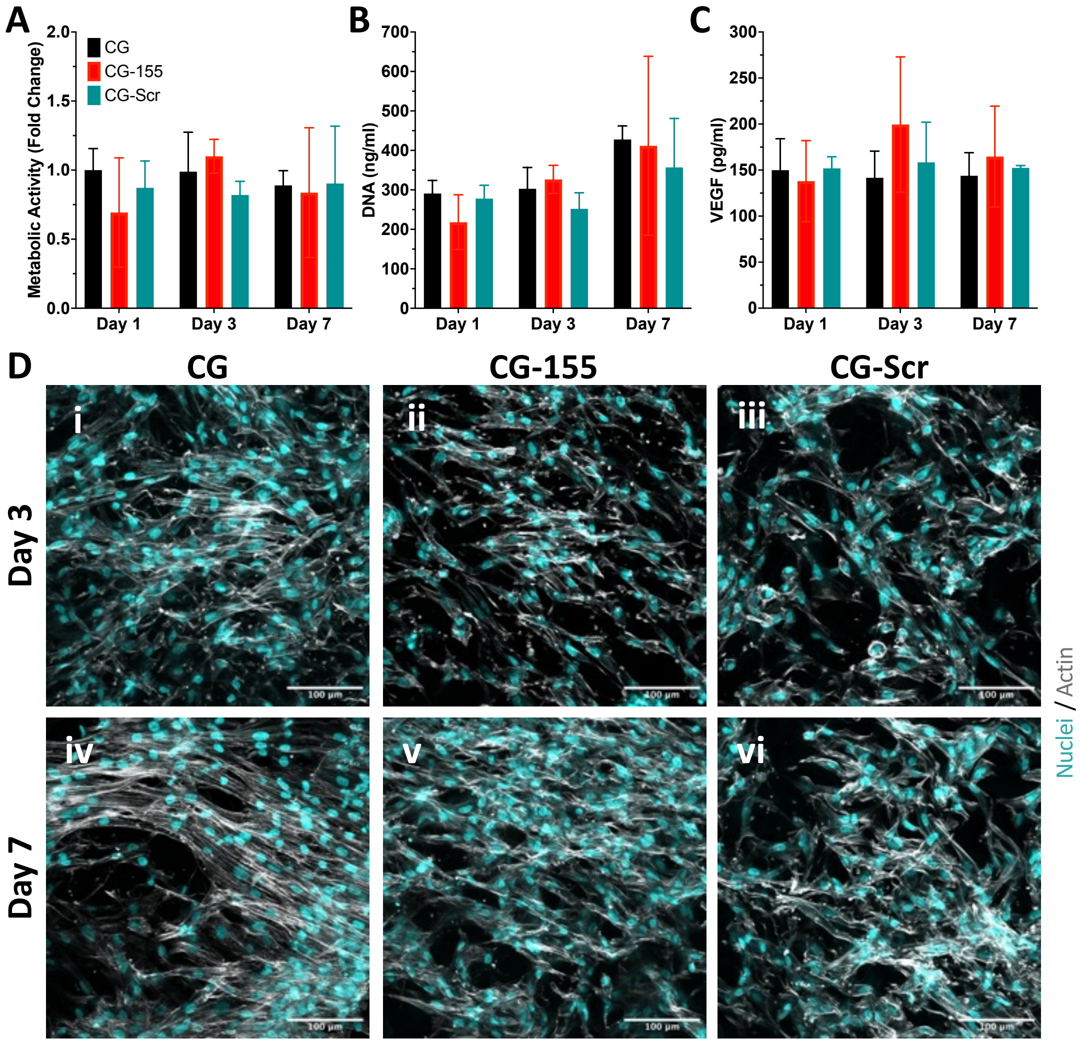


**Supplementary Figure 3. CG-155-i scaffolds do not elicit a detrimental response in human dermal fibroblasts**. A-B) Assessment of cell viability through metabolic activity and DNA quantification does not show any significant difference among groups. C) Scaffold-mediated VEGF expression from HDFs does not reveal clear variation for up to 7 days post-transfection. D) Visualisation of cellular morphology displays the distinctive spindle-like morphology of fibroblasts on all scaffold groups. Data shows mean ± SD.

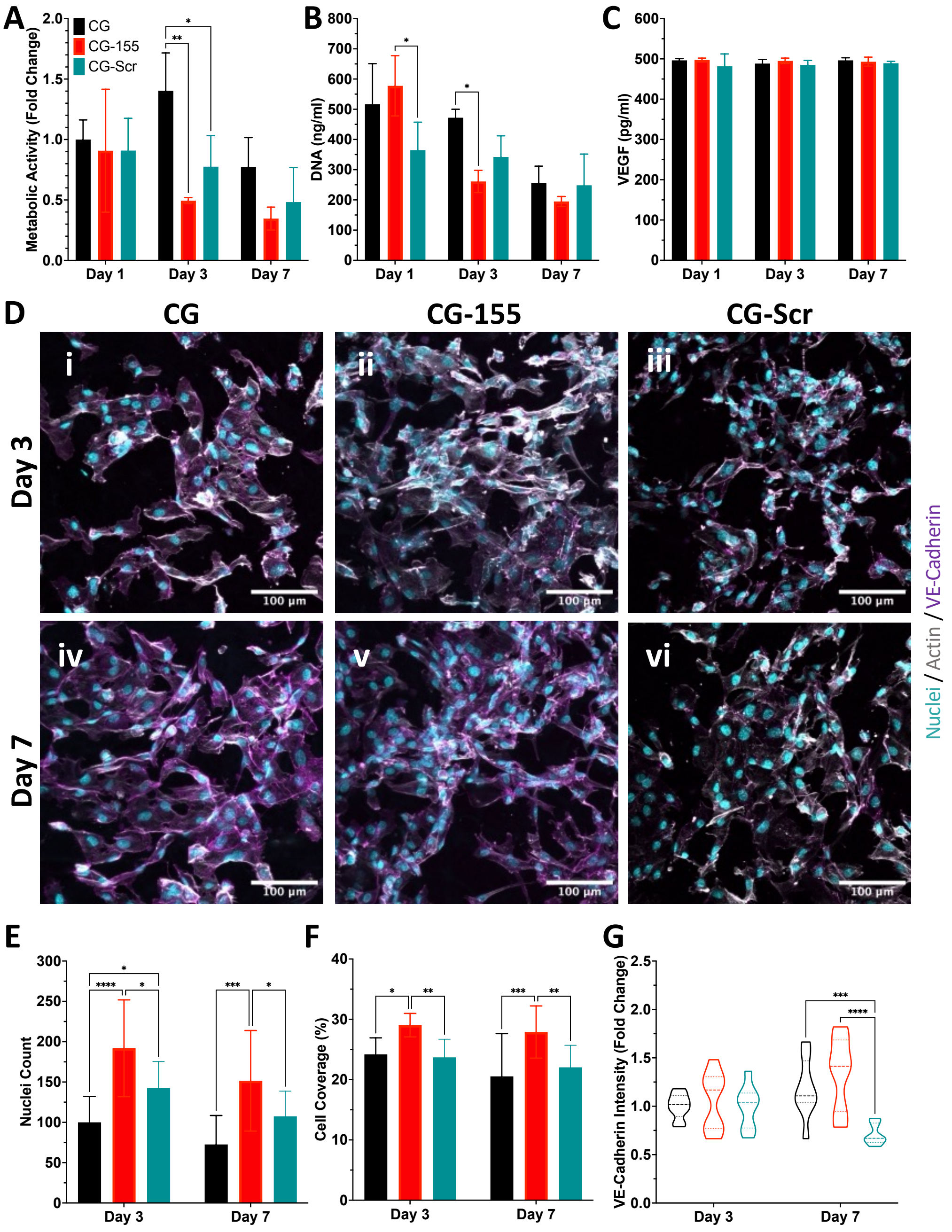


**Supplementary Figure 4. CG-155-i scaffolds support endothelial cell proliferation and the expression of the key endothelial cell surface marker VE-cadherin**. A-B) Analysis of cell viability through metabolic activity and DNA quantification exhibits decreased HUVEC viability on day 3 post-transfection on CG-155-i scaffolds. C) Assessment of VEGF expression does not display any clear differences among scaffold groups. D) Imaging of HUVECs on scaffolds shows a diminished VE-cadherin intensity from HUVECs on CG-Scr-i scaffolds on day 7 post-transfection. E-G) Nuclei count, cell coverage, and VE-Cadherin intensity quantification reveal an improved response from HUVECs on CG-155-i scaffolds while supporting the observation of reduced VE-cadherin expression on CG-Scr-i scaffolds. Data shows mean ± SD, * indicates $p<0.05$, ** $p<0.01$, *** $p<0.001$, **** $p<0.0001$.

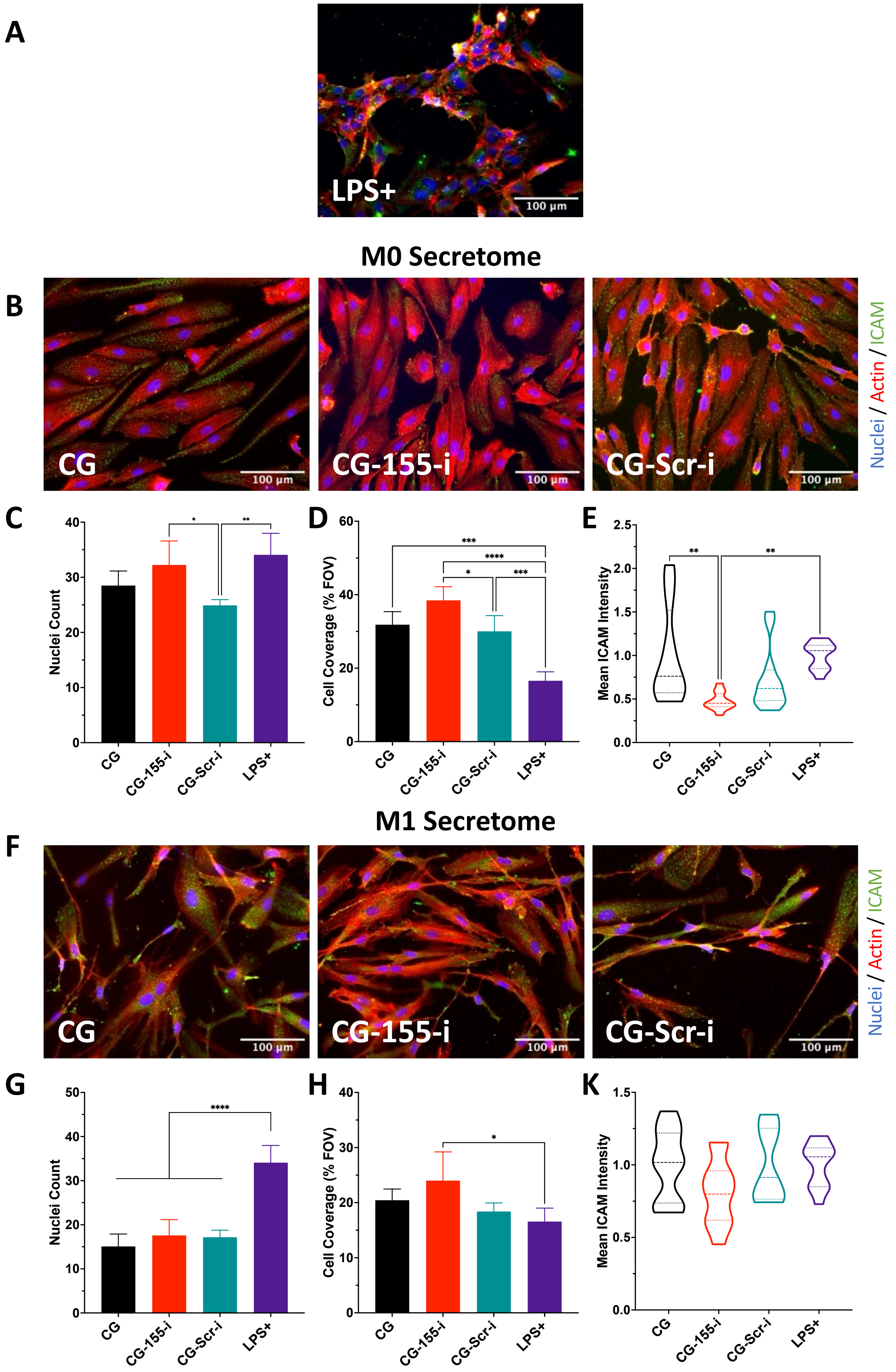


**Supplementary Figure 5. Secretome from macrophages on CG-155-i scaffolds induces anti-inflammatory responses on endothelial cells.** A) Representative image of LPS-treated endothelial cells used as positive control.B-E) Endothelial cells exposed to M0 macrophages secretome show a reduced expression of the pro-inflammatory marker ICAM in the CG-155-i group. F-K) The influence of M1 macrophage secretome on endothelial cells exhibits clear morphological changes and decreased ICAM intensity. Data shows mean ± SD, * indicates p<0.05, ** p<0.01, *** p<0.001, **** p<0.0001.

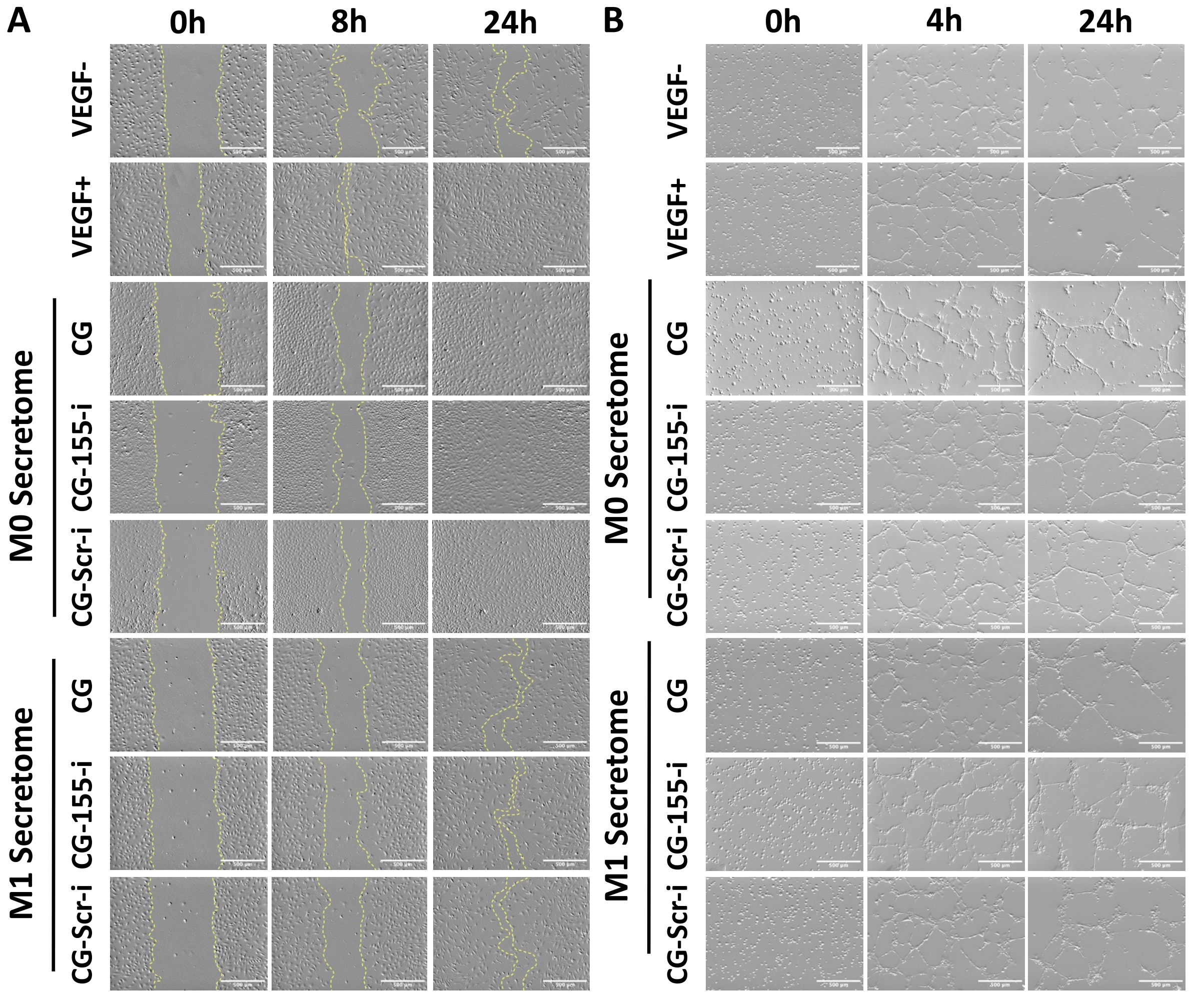


**Supplementary Figure 6. Secretome from macrophages on CG-155-i scaffolds induces faster endothelial cell migration and organisation.** A) Representative images of endothelial cells used for assessment of migration rate following exposure to secretome from macrophages on miRNA-i-activated scaffolds. B) Representative images of endothelial cells used for assessment of vascular-like organisation following exposure to secretome from macrophages on miRNA-i-activated scaffolds. Scale bar = 500 um
